# Collaborative Augmented Reconstruction for Scaled Production of 3D Neuron Morphology in Mouse and Human Brains

**DOI:** 10.1101/2023.10.06.561172

**Authors:** Lingli Zhang, Lei Huang, Zexin Yuan, Yuning Hang, Ying Zeng, Kaixiang Li, Lijun Wang, Haoyu Zeng, Xin Chen, Hairuo Zhang, Jiaqi Xi, Danni Chen, Ziqin Gao, Longxin Le, Jie Chen, Wen Ye, Lijuan Liu, Yimin Wang, Hanchuan Peng

## Abstract

Digital reconstruction of the intricate 3D morphology of individual neurons from microscopic images is widely recognized as a crucial challenge in both individual research laboratories and large-scale scientific projects focusing on cell types and brain anatomy. This task often fails both conventional manual reconstruction and state-of-the-art automatic reconstruction algorithms, even many of which are based on artificial intelligence (AI). It is also critical but challenging to organize multiple neuroanatomists to produce and cross-validate biologically relevant and agreeable reconstructions in scaled data production. Here we propose an approach based on collaborative human intelligence augmented by AI. Specifically, we have developed a Collaborative Augmented Reconstruction (CAR) platform for neuron reconstruction at scale. This platform allows for immersive interaction and efficient collaborative-editing of neuron anatomy using a variety of client devices, such as desktop workstations, virtual reality headsets, and mobile phones, enabling users to contribute anytime and anywhere and take advantage of several AI-based automation tools. We have tested CAR’s applicability for challenging mouse and human neurons towards a scaled and faithful data production. Our data indicate that the CAR platform is suitable for generating tens of thousands of neuronal reconstructions used in our companion studies.

## Introduction

Three-dimensional (3-D) neuron morphometry offers direct insights into the complex structures and functions of individual neurons and their networks, enhancing our understanding of a brain and its capabilities (BICCN, 2021; Hawrylycz, et al., 2023). The pursuit of detailed morphometric measurements of neurons, particularly at the single-cell level and throughout an entire brain, has garnered several seminal datasets including several thousand fully reconstructed neurons in mouse brains (Winnubst, et al., 2019; Peng, et al., 2021; Gao, et al., 2022). The generation of these morphology datasets became possible thanks to both recent advancements in sparse neuron labeling and light microscopy imaging of whole brains (e.g. Rotolo, et al., 2008; Kuramoto, et al., 2009; Ghosh, et al., 2011; Gong, et al., 2016; Lin, et al., 2018; Matho, et al., 2021; Munoz-Castaneda, et al., 2021; Wang, et al., 2021; Han, et al., 2022), and particularly new development of neuron reconstruction (also called neuron tracing) technologies from 3-D light microscopy images (Liu, et al., 2022; Manubens-Gil, et al., 2023).

A common goal of neuron reconstruction methods is to reconstruct digital models of the complete neuronal morphology with a low error-rate (Garvey, et al., 1973; Peng, et al., 2011; Acciai, et al., 2016; Peng, et al., 2017; Liu, et al., 2022; Manubens-Gil, et al., 2023). As neuron tracing techniques can be intuitively categorized as manual, semi-automatic, and automatic methods, it is obvious that varying levels of automation enabled by computer algorithms and the manual involvement of human labors have a deterministic impact on the efficiency and productivity of the digital reconstruction. Despite a number of claimed successes in automated neuron tracing, practically the majority of automation has only been applied to fairly simple use-cases where the signal-to-noise ratio is high, or the entirety of neurite-signal is not required to be traced (Liu, et al., 2022). Indeed, as the worldwide community has recognized that there exists no single best algorithm for all possible light-microscopy neuronal images (Peng, Hawrylycz, et al., 2015; Peng, Meijering, et al., 2015), careful evaluation of automated tracings must be cross-validated before they may acclaim biological relevance (Manubens-Gil, et al., 2023). Therefore, a remarkable, unanswered key question in the field is how to produce 3-D reconstructions of complicated neuron morphology at scale, while ensuring these reconstructions are both neuroanatomically accurate and reliable. This is the central question we address in this study.

We believe that the ultimate achievement of large-scale neuron-morphology production will entail harnessing automation algorithms and increasingly powerful computing hardware to augment data production rates within specified timeframes. To reach such a goal, we considered practical challenges that must be surmounted. Neuron morphology encompasses a multitude of delicate characteristics, including the presence of thin yet extensive neurite fibers and spines, as well as the intricate broken signal patterns along neurites caused by the uneven distribution of light indicators (e.g. fluorescent proteins) during the neuron labeling process (Peng, et al., 2011). It is imperative to exercise caution to prevent unintentional compromise of these structures throughout tracing and preliminary processing steps, such as image preprocessing (Li, et al., 2017; Klinghoffer, et al., 2020; Jiang, et al., 2021; Zhang, et al., 2022). In addition, neurons frequently possess complex structures that can hinder the attainment of unequivocal observations. This complexity can become magnified when a region contains multiple neurons, and very large projecting neurons need to be reconstructed from whole brain images which contain trillions of voxels. Due to these hurdles, currently there remains a substantial scarcity of high-quality training datasets of neuron morphology, making the development of deep-learning and similar machine-learning methods for this task a formidable challenge (Liu, et al., 2022). In this context, the recent achievements in the series of generative pre-trained transformers (GPT) (Open AI, 2023) have opened promising avenues for the development of Large Language Models (LLM) (Brown, et al., 2020; Zhao, et al., 2023) and the pursuit of Artificial General Intelligence (AGI) (Goertzel, et al., 2007). Meanwhile, there have been concerted efforts to develop general-purpose models for computer vision tasks (Radford, et al., 2021; Kirillov, et al., 2023), including extended applications in the domain of medical image processing (Mazurowski, et al., 2023). However, it is imperative to exercise caution when employing such tools to prevent potential controversies and common pitfalls (Bubeck, et al., 2023). Furthermore, it is worth noting that their capabilities are currently substantially limited when it comes to annotating large-scale multidimensional imaging data (He, et al., 2023).

While the challenges in neuron reconstruction are significant and cannot yet be fully addressed through pure artificial intelligence (AI) approaches, we have taken a proactive step towards overcoming these hurdles. We develop a comprehensive technical platform, called Collaborative Augmented Reconstruction (CAR), to enable many annotators and end-users to contribute annotating very complicated 3-D morphology in a collaborative way, which at the same time is also enhanced to generate neuroanatomically plausible reconstructions by leveraging specifically designed AI tools to automate the data production for critical neuron skeletons and completeness of neuron morphology. In this way, we have effectively integrated collective intelligence with artificial intelligence for the task of neuron reconstruction from large-scale 3D brain images, resulting in a crowd-in-the-loop design that boost both the biological significance of the reconstructed morphology and the production speed.

In this manuscript, we showcase CAR’s effectiveness in several applications for challenging mouse and human neurons towards a scaled and faithful data production. Our data indicate that the CAR platform is suitable for generating tens of thousands of neuronal reconstructions used in our companion studies. We have adopted CAR as a major morphological data generation platform in several ongoing projects including The BRAIN Initiative Cell Census Network (BICCN) and BigNeuron (Manubens-Gil, et al., 2023).

## Results

### The Collaborative Augmented Reconstruction platform enables versatile morphometry in real-time

Our major result in this study is to develop a Collaborative Augmented Reconstruction (CAR) platform for the complex challenges of 3-D neuron reconstruction from noisy, large 3-D light microscopy images of mammalian brains. Compared to other neuron reconstruction software packages, CAR stands out as a versatile computational platform (**Supplementary Table 1,2**). It was specifically designed to address the challenges associated with faithful reconstruction of neuronal images in mammalian brains, with a particular emphasis on the mouse and human brains. CAR’s scalability also allows it to cater to a wide range of neuroscience applications (**Fig. 1**). One key strength is its accessibility across various client-end devices with built-in artificial intelligence (AI) components, including regular desktop/laptop computers, virtual reality headsets, mobile phones, and even game consoles. This extensive device compatibility enables efficient visualization and confident annotation of intricate 3D neuroscience data (**Fig. 1A,B, Supplementary Fig. 1**). These diverse client options offer specific advantages for neurodata validation by providing evidence of its completeness and accuracy. With the CAR platform, we successfully organized a geographically dispersed team to collaborate effectively, as shown in several applications later in the paper. Team members worked together in real-time within a shared virtual environment, allowing them to view and interact with each other’s annotations promptly, with the assistance of real-time AI-powered tools. Furthermore, CAR offers the flexibility for users to work independently while maintaining data synchronization among the team, fostering seamless collaboration.

**Fig. 1.**
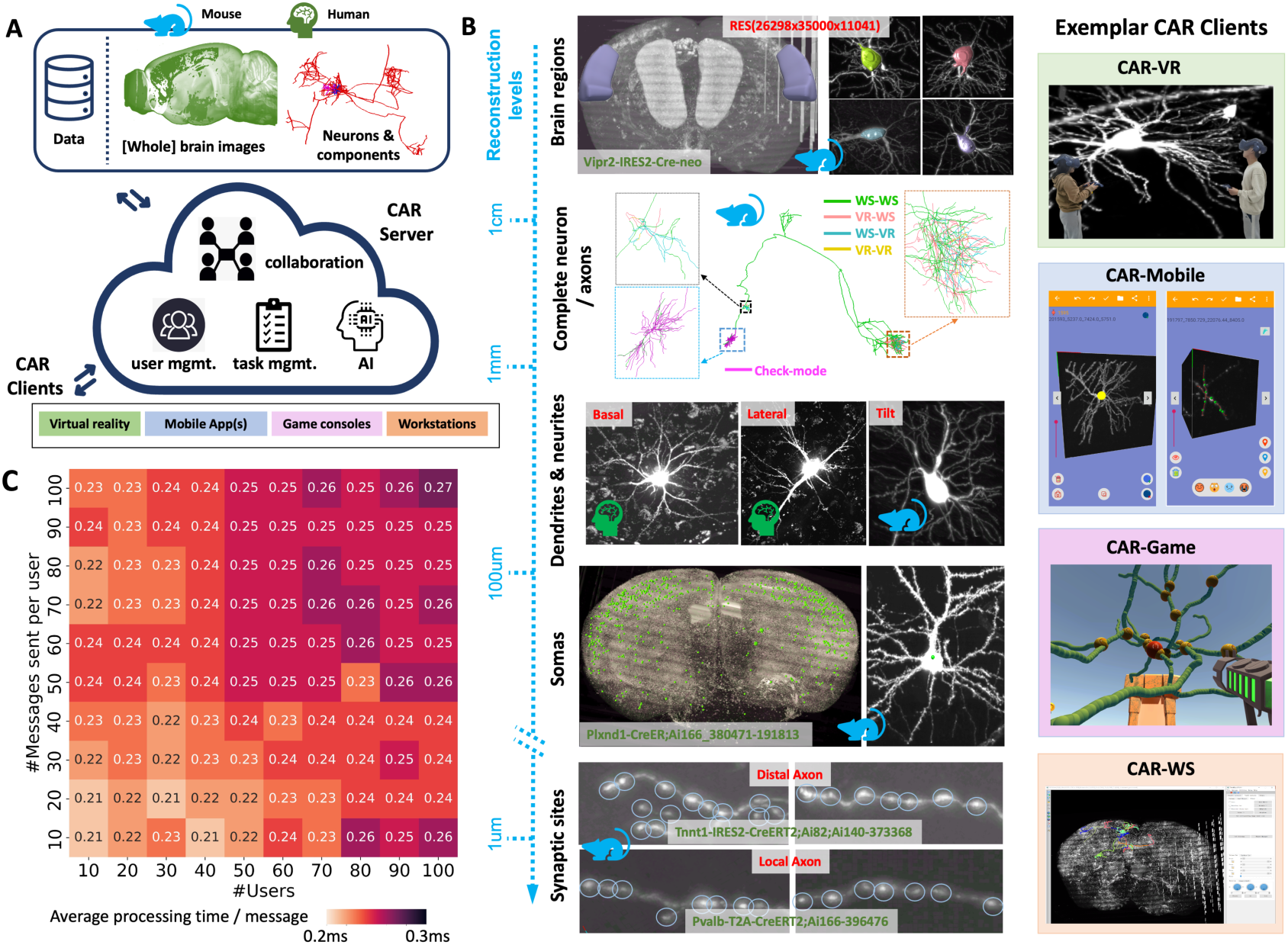
The CAR platform and its neuroscientific applications. **A,** The framework of the CAR platform. CAR has a cloud-based architecture and supports diverse types of clients, including workstations, virtual reality tools, game consoles and mobile Apps. CAR is built upon a comprehensive collaboration mechanism, provides management for data, tasks, and users, and is boosted by artificial intelligence capabilities. The system addresses the challenges regarding faithful neuron reconstruction, with a particular focus on mouse and human brain data. **B,** Left: Typical morphometric tasks performed using CAR at different reconstruction levels from centimeter-to micrometer-scales, e.g., brain regions location tagging, complete neurons reconstruction (WS-VR: neurites annotated by workstation (WS) and validated by virtual reality (VR); WS-WS, VR-WS, VR-VR: similar meanings; check-mode: neurites that are reconstructed yet validated), neurite tracing, soma annotation, bouton detection and proofreading. Right: Exemplar CAR clients are showcased, including CAR-VR (also called TeraVR+), CAR-Mobile (also called Hi5), CAR-Game (also called BrainKiller, unpublished work), and CAR-WS (also called TeraFly-x). **C,** The performance of the CAR server under highly concurrent scenarios. In CAR, the annotation operations were synchronized among the server and the users using network messages. We modeled situations where many collaborating users (ranging from 10 to 100) were sending a burst number of messages. The heatmap shows the average processing time at the CAR server for each message. The y-axis indicates the number of messages sent per user, while the x-axis represents the number of users engaged in concurrent tasks.

We utilized CAR to investigate brain anatomy across various scales. Specific tasks include tagging 3D brain regions, reconstructing entire neurons, tracing local dendritic and axon arbors, identifying somas, verifying potential synaptic sites, and making various morphometric measures (**Fig. 1B**). These tasks often necessitated collaboration among team members who utilized different types of CAR clients. For instance, in a shown example (**Supplementary Fig. 2**), five users used CAR and collaborated to reconstruct a complete neuron from whole-mouse-brain imaging data, with different clients focusing on reconstruction of terminal branches, middle branches, and bifurcation or a crossing, respectively. Furthermore, game consoles were employed to validate the topological accuracy of the reconstruction. As CAR benefits a team from enhanced productivity and communication, CAR facilitates improved comprehension of complex neuron structures and knowledge sharing among users who might be geographically dispersed.

CAR’s cloud server manages centralizing operations, synchronizing annotation data, and resolving any conflicts that may arise (**Fig. 1A**). All data, including 3D microscopic brain images and reconstructed neuron morphology, are hosted in the cloud storage, so users do not need to maintain them locally at CAR clients. We found that the CAR server was capable of handling large numbers of users and message-streams in real-time. Indeed, the CAR server responded within 0.27ms even for 10,000 concurrent messages (**Fig. 1C**).

### CAR reconstructs whole-brain projection neurons with converging correctness

We tested CAR on challenging 3-D annotation tasks that encompassed large, multi-dimensional, and innovative datasets. In the first application, we applied CAR to annotating complicated 3-D morphologies of large projection neurons in whole mouse brains, where a typical testing dataset involves an XYZ volumetric brain image with about 40,000×30,000×10,000 voxels, or 12 tera-voxels. CAR allows annotating an initial neuron morphology-reconstruction that has been generated using either an automatic neuron-tracing algorithm or from scratch (**Methods**). The accomplishment of the large-scale reconstruction is ensured through a series of CAR components, including CAR-WS and CAR-VR for instance that have robust large-data handling capabilities.

We focused on representative neuron types in the mouse brain, with their cell bodies situated in 20 anatomical regions corresponding to major functional areas, including cortex, thalamus, and striatum (**Fig. 2A**). These neurons form a broad coverage in the brain with often remarkably long axons (**Fig. 2A - left panel**). They also have dramatically variable 3-D morphology in terms of projection target-areas, projection length (about 1.90 cm to 11.19 cm) and complexity in their arbors (with about 300 to 1300 bifurcations) (**Fig. 2A - right panel**). With the aid of CAR, we achieved reconstruction accuracy of over 90% for all test neurons (**Fig. 2A**), accomplished with the collaborative efforts of layman-annotators, and validated by additional expert-gatekeepers.

**Fig. 2.**
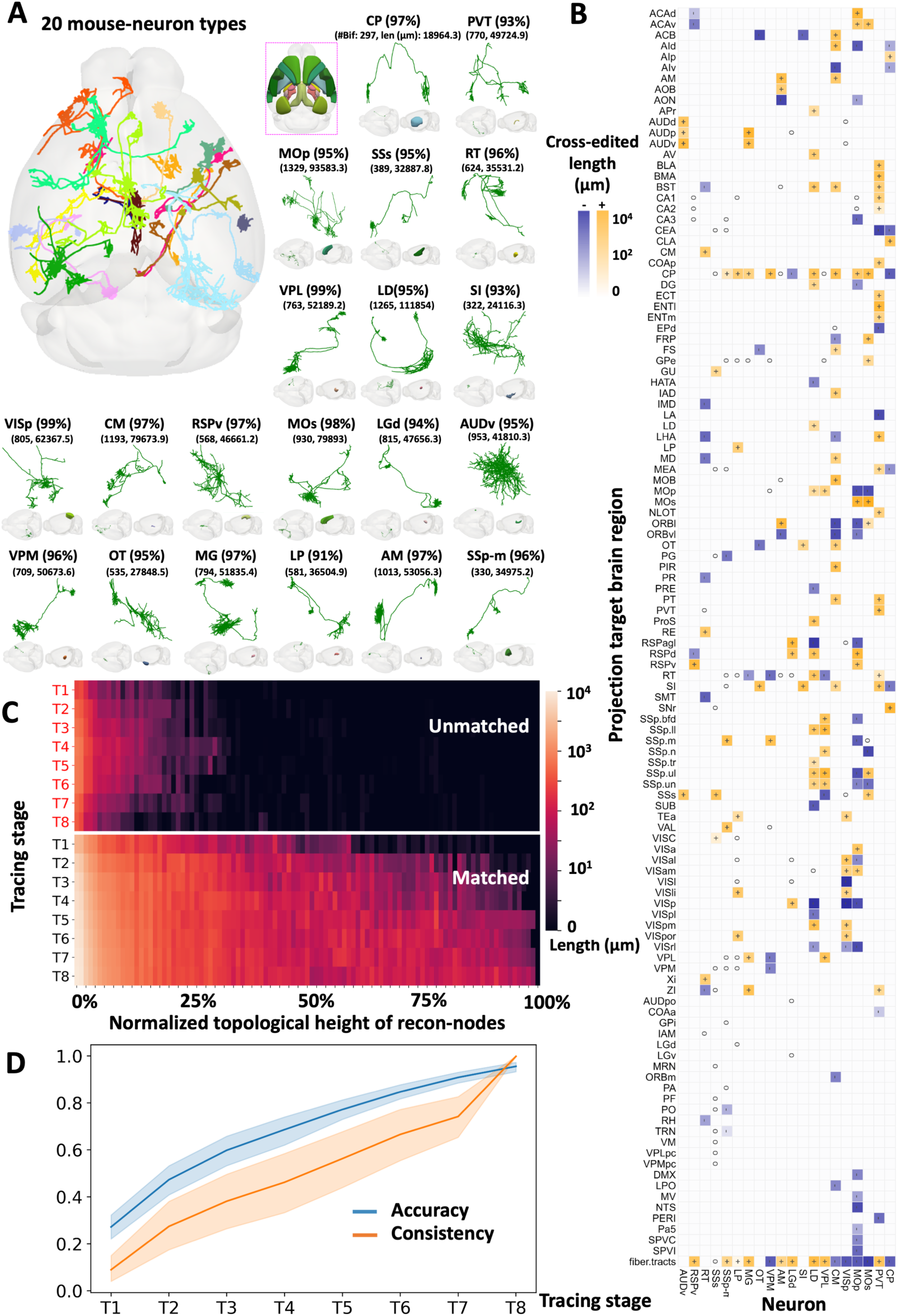
Reconstruction of complete mouse neurons using CAR. **A,** The complete reconstruction of exemplar mouse-neurons from 20 different brain regions. Left: The visualization of a top-down view of the exemplar neurons registered to the standard Allen brain atlas. Each color represents an individual neuron, and the small panel on the right indicates the respective brain region to which these neurons belong. These 20 different neurons are registered to CCFv3 using mBrainAligner (Qu, et al., 2022). Right: Visualization of the neurons separately, providing their type, reconstruction accuracy, number of bifurcations and total length (μm) (calculated using Vaa3D plugin “global feature”). The mapped morphology in the standard atlas and the brain region that the neuron origins are also visualized below each neuron. **B,** The projection map illustrates the lengths of reconstructed neurites contributed through collaborations. The horizontal and the vertical axis represent the origin (soma location) and destination (projection location) regions, respectively. Each cell in the map represents a projection pair, with the darkness of the shade corresponding to the amount of the cross-edited length by collaboration. A ’+’ symbol (yellow) is employed to denote cases where collaborative-addition was the predominant operation, while a ’-’ symbol (purple) is used for instances where collaborative-subtraction dominated the editing process. The symbol ’o’ indicates that no editing was performed through collaboration. **C,** The normalized topological height of recon-nodes (**Methods**) of matched and unmatched structures for 20 neurons. This plot compares the topological height of reconstructed nodes with expert results at eight stages along the reconstruction timeline. Matched structures (bottom panel) indicate successful reconstructions that align with expert results, while unmatched structures (upper panel) deviate from expert results. **D,** Average accuracy and user consistency in 20 neurons across eight tracing stages. Blue polyline: average accuracy (blue shade: 95% confidence interval); orange polyline: consistency (**Methods**) among the contributors (orange shade: 95% confidence interval).

As shown in an example of synergized annotation of a VPL neuron done by six layman contributors, where all neurite-segments were cross-validated (**Supplementary Fig. 3,4,5**), an additional expert neuroanatomist further examined the reconstruction and adjusted only less than 2% of the neuron’s substructures in this case (**Supplementary Fig. 4C**), which indicates that the laymen’s consensus was comparable with the expert. When we visualized the heatmap of user participation in editing different regions of the neuron, it showed the most intensive user collaboration happened around dendrites and distal axonal clusters that correspond to complicated branching structures (subpanels of **Supplementary Fig. 4D**).

Since the projecting targets of neurons hold essential information about their roles within the brain, we compared the projection maps derived from collaborative reconstructions and non-collaborative reconstructions performed by the same group of annotators. Through collaboration, we achieved a total neurite length of 84.8cm for the 20 neurons. We also created a contrast map illustrating the edited differences between these two versions (**Fig. 2B**), revealing a total variation (including both additions and subtractions) in neurite length amounting to 37.3cm. In other words, nearly 44% of the structures of these projection neurons underwent significant cross-editing (**Supplementary Fig. 6**). Notably, the non-collaborative version exhibited numerous instances of erroneously connected or missing neurites, which could considerably undermine subsequent analyses. In this context, the ability to cross-validate the reconstructions of projection neurons, as facilitated by the collaborative annotation approach of CAR, becomes crucial.

A notable advantage of employing CAR is its capacity to identify potential unmatched (incorrect) reconstructions and avert unfavorable consequences. In other words, this platform possesses the ability to limit potential errors and progressively refine the reconstruction process until a consensus is achieved among contributors. To facilitate quantitative analysis across different neurons, we defined a “normalized topological height” (NTH) for reconstruction nodes within a neuron (**Supplementary Fig. 7; Methods**). NTH indicates the corrective effort required to rectify a reconstruction error involving a particular node and all its subsequent branching structures (**Methods**). The magnitude of the height directly correlates with the cost of modification. Across all tested mouse neurons, we observed a gradual reduction in the proportion of incorrect reconstruction components over both the tracing stage and the NTH (**Fig. 2C, Supplementary Fig. 8**). Notably, these errors remained confined to regions with low topological heights, suggesting that most reconstruction inaccuracies were promptly rectified before they could give rise to further erroneous structures. In this way, CAR excels in both reconstruction accuracy and efficiency.

Finally, we observed a consistent enhancement in overall reconstruction accuracy toward greater than 90% as agreement among contributors steadily increased over time (**Fig. 2D**). CAR facilitates such beneficial, intertwined collaboration, allowing each user to actively review other contributors’ reconstructions while simultaneously receiving assistance from fellow users.

### AI-based branch and terminal classifiers for automating reconstructions

One key feature of CAR is to augment the throughput of neuron reconstruction using two AI tools based on Convolutional Neural Networks (CNN) (**Fig. 3, Supplementary Fig. 9**). First, a Branching Point Verifier (BPV) was developed to determine if the branching points in a reconstruction correspond to real bifurcation loci in the imaging data (**Supplementary Fig. 9A**). BPV combines the advantages of attention mechanism and residual blocks to efficiently extract distinctive neuronal image features (**Methods**). Second, a Breakpoint Detector (BD) was designed to identify potential interruption in tracing neurites by classifying real neurite terminals against potential early termination in tracing (**Supplementary Fig. 9B**). In order to better distinguish endpoints and breakpoints whose image share similar features, BD allows the network to learn more distinctive features (**Methods**). Both BD and BPV were deployed at the CAR cloud server to periodically assess the neuron reconstructions, followed by pushing various suggestions of potentially erroneous breakpoints and branching points to CAR clients. This AI-augmented interactive annotation was effective. Indeed, BD and BPV behave like independent AI-collaborators (contributors), frequently reminding human users to fix mistakenly reconstructed branching structures and continue tracing from forgotten breakpoints (**Fig. 3A**).

**Fig. 3.**
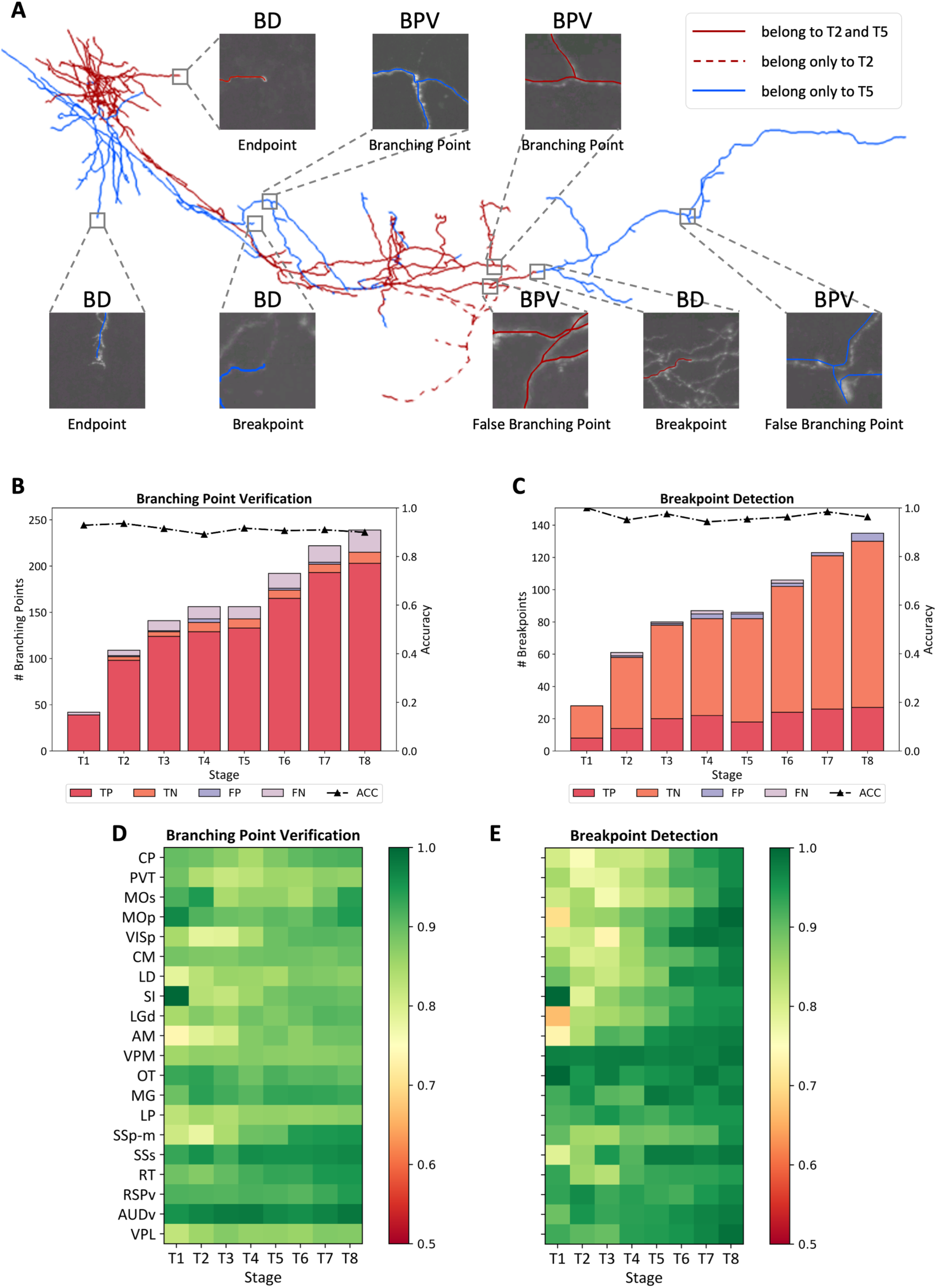
Collaboration of AI agents in CAR during reconstruction stages T1 ∼ T8. **A,** A half-way reconstructed OT neuron. The red colored neurites (both in solid and dashed lines) comprised the morphology at T2, while the neurites shown in solid lines (both in red and blue) formed the morphology at T5. Some incorrectly reconstructed parts in T2 were shown in dashed lines, and were deleted by the time of T5, thanks to the hints provided by BPV. The subpanels showed the positive and negative cases of BPV and BD, together with the image at the local region. **B, C** The performance of BPV and BD at 8 stages, respectively. For each stage, the number of true positive (TP), true negative (TN), false positive (FP), false negative (FN) samples were plotted, as well as the accuracy (**Methods**). **D, E,** The accuracy of the two models for all the 20 neurons at 8 stages. Horizontal axis: stage; vertical axis: neuron type; color map: accuracy.

For instance, for the OT neuron (**Fig. 2A**), compared to the expert-validated reconstruction, the overall accuracies of both BPV and BD were above 90% throughout the reconstruction process (**Fig. 3B, C**), even for partially completed neuron reconstructions. We also tested the reliability of both tools for other types of projection neurons that have many thin, often broken axonal branches (**Fig. 2A**). We observed that during the entire reconstruction process, BD and BPV constantly yielded an average accuracy over 90% and 85%, respectively (**Fig. 3D, E**). This means that our AI tools were reliable to produce faithful hints for human curation, largely independent of the completeness of reconstructions.

### Reconstruct human cortical dendrites in 10 brain regions

In our effort to trace complete mouse neurons with long, faint projecting axons (**Fig. 2,3**) that are labeled using genetic and viral methods (Daigle, et al., 2010, Gong, et al., 2016), the dendrite-tracing parts of those neurons are not as hard as axon-tracing because of the relatively low noise level in the respective dendritic areas in the mouse brain images. Here, we have also applied CAR to reconstruct challenging human cortical neurons where their dendritic images have abundant noises, due to various artifacts of dye-injection, which is another widely used method for neuron labeling.

We specifically considered human cortical neurons being routinely generated in a consortium involving human-neuron extraction, labeling, mapping, reconstruction, and modeling using a human Adaptive Cell Tomography method (Han, et al., 2022). While human brain images can be obtained in high-throughput through perfusion and imaging, the noise level is substantial, because of the fluorescence of blood vessels and leaked dye out of injected cell bodies or other injection sites. These human neuron images often fail other neuron tracing methods. We used CAR to reconstruct 80 human neurons from 10 cortical regions (**Fig. 4A -column 2 of the array, Supplementary Fig. 10**). These neurons are mainly pyramidal cells with around 100 branches and 15∼20 topological layers of bifurcations embedded in images with intense noises (**Fig. 4A -columns 1,5 of the array, Fig. 4B**). The reconstruction results showed that annotators effectively collaborated on reconstructing various parts of these neurons (**Fig. 4A - column 3 of the array**), especially focusing on areas with high branching density where the structural complexity was large (**Fig. 4A - columns 4 of the array**).

**Fig. 4.**
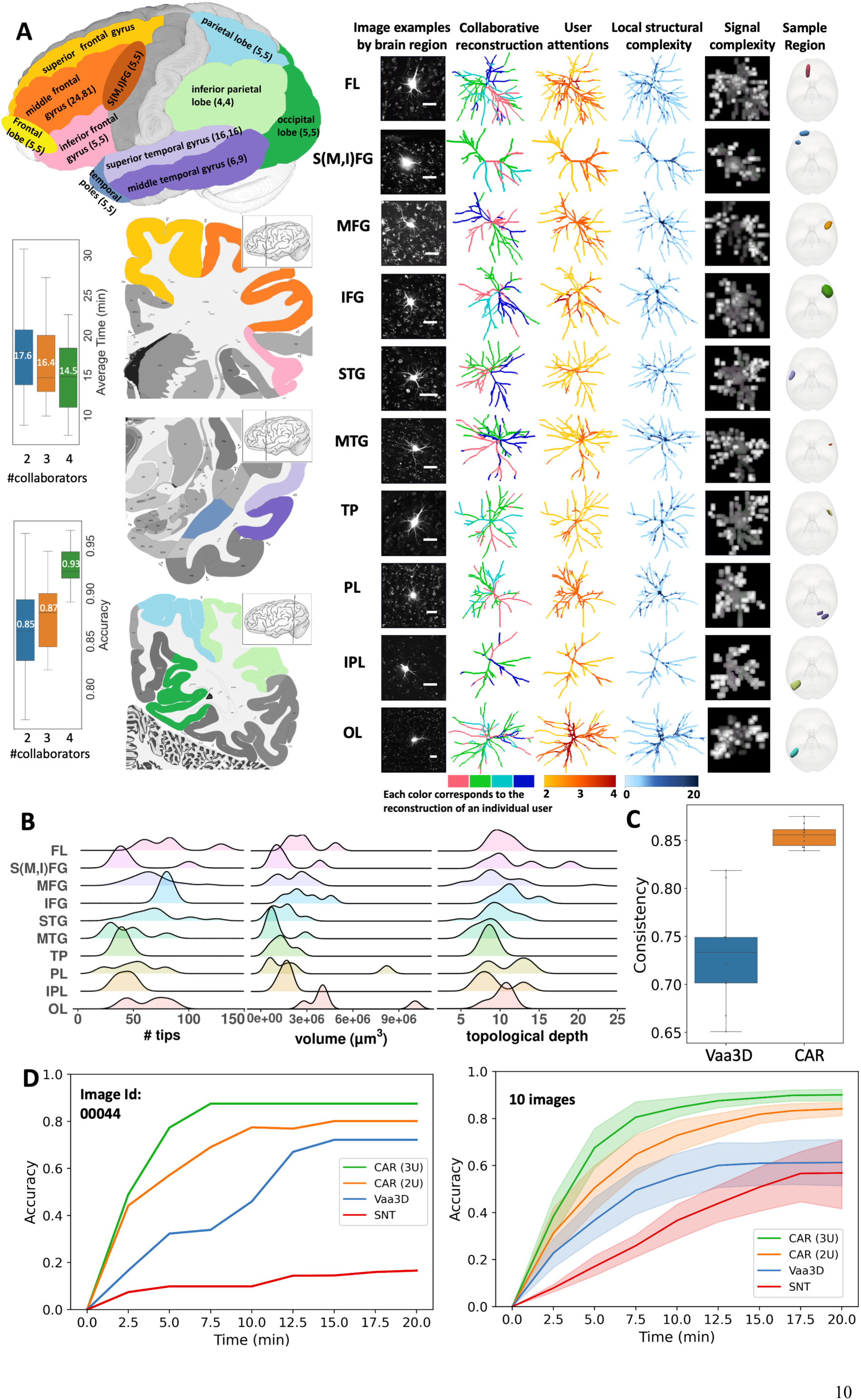
CAR used in human neuron reconstruction. **A,** Left: Semantic mapping scheme of human brain regions reconstructed. Each brain region (see **Methods** for region names) is represented by a different color. For each region, the two numbers indicate the number of individual neurons reconstructed and the total reconstruction copies (some neurons were reconstructed more than once for validation), respectively. Right: Representative cells for different cell types are displayed in each row. From left to right, each column gives the images (scale bar = 50 μm), user participations (each color corresponds to the annotation of an individual user), user attention heatmaps (color indicates the number of editing users for the region) (**Methods**), local structural complexity (color indicates the complexity of the local structure) (**Methods**), imaging signal complexity (intensity indicates the signal complexity in the local region) (**Methods**), the actual locations where brain tissues were extracted respectively, each represented by a different color. These extracted regions were then mapped and overlaid onto the standard brain coordinate template MNI152. The boxplot on the left illustrates the relationship between reconstruction time and achieved accuracy for different collaboration patterns across all neurons (Numbers in white color: average). **B,** The number of tips, the volume, and the topological depth for the neurons in the 10 types. These three features are defined in L-Measure (Scorcioni, et al., 2008). **C,** The consistency of reconstructions of different Vaa3D trials and CAR trials. The Vaa3D plugin “neuron distance” was used to calculate the consistency measure. **D,** Comparison of the reconstruction accuracy over time among different tools: CAR with 3 users, CAR with 2 users, Vaa3D-3Dviewer with 1 user and SNT with 1 user. On average, CAR (3U) achieved convergence within approximately 10 minutes. To enhance comparability, a 20-minute upper time limit was applied to all tools tested, which is twice the convergence time of CAR (3U). The left panel showcases an example neuron (ImageID: 00044), while the right panel displays all the 10 neurons (shades: 95% confidence interval).

It was interesting to observe that as the number of collaborators using CAR increased from 2 to 4, neurons were reconstructed with 7% to 18% less time, while the overall error reduced from above 15% to as little as 7% steadily (**Fig. 4A**). The collaboration of four contributors showed promise in reconstructing 15 randomly selected neurons with varying signal-to-noise ratios. Their combined effort yielded a dependable accuracy rate of approximately 91% (**Supplementary Fig. 11**).

Thanks to its built-in immersive visualization capability and the collective consensus among annotators, CAR can generate more stable results than alternative approaches that do not optimize collaborative reconstructions. For instance, the inter-group consistencies in using CAR and Vaa3D’s 3D-Viewer (Peng, et al., 2014a) were 0.86 and 0.73, respectively, when two groups of annotators were tasked with completing the same reconstruction (**Fig. 4C; Methods**). The advantage of CAR over conventional tools becomes even more pronounced when testing is performed on human neuron images with a high noise level. In the case of a randomly chosen testing image (#00044), CAR achieved an 85% accuracy of reconstruction within 7.5 minutes, whereas the tool Vaa3D’s 3D-viewer required 15 minutes but yielded inferior results (approximately 70% accuracy). Similarly, a 2-D visualization-based tool, SNT (Arshadi, et al., 2021), needed 20 minutes but still missed over 80% of neurites (**Fig. 4D - left panel**). In a comparison involving 10 randomly selected neurons and expert-validated reconstructions, a consistent pattern emerged: CAR began converging to over 80% accuracy in about 7.5 minutes, whereas the respective Vaa3D and SNT achieved a maximum of 50% to 60% accuracy at the expense of nearly double or triple the reconstruction time (**Fig. 4D - right panel**).

### Reconstruct somas and synaptic sites at whole-brain scale

In addition to generating reconstructions of complex axons and dendrites toward full neuron morphology as shown above, we also applied CAR to produce other types of digital reconstructions involving sub-structures of neurons at the whole brain scale. One illustrative example is our application of CAR to detect somas in mouse brains. To do so, we first employed an automatic soma detection algorithm (**Methods**) to identify potential soma locations across diverse mouse brains, and then used the CAR cloud server to dispatch image-blocks containing the putative somas to many remote users who use CAR’s mobile client (**Fig. 1**). These users were able to fine-tune the soma locations in real-time, cross-validated the results, and typically completed annotating each image-block within a few seconds.

By employing this protocol, we generated one of the most extensive databases of annotated somas, utilizing genetically labeled neurons across 58 whole mouse brains, which spanned a comprehensive total of 609 brain regions, all aligned to the Allen Common Coordinate Framework (**Fig. 5A**). Specifically, we used CAR-Mobile client to accurately identify 156,190 somas, within approximately 4 weeks, involving collaboration among 30 users (23 trained users and 7 novice annotators) (**Fig. 5A**). Given the heightened precision of soma locations validated through CAR-Mobile client compared to the initial automated detection, we were able to proceed with the further reconstruction of complicated neuronal morphologies within specific brain regions, such as hippocampus and striatum (**Fig. 5A**), still based on the CAR platform.

**Fig. 5.**
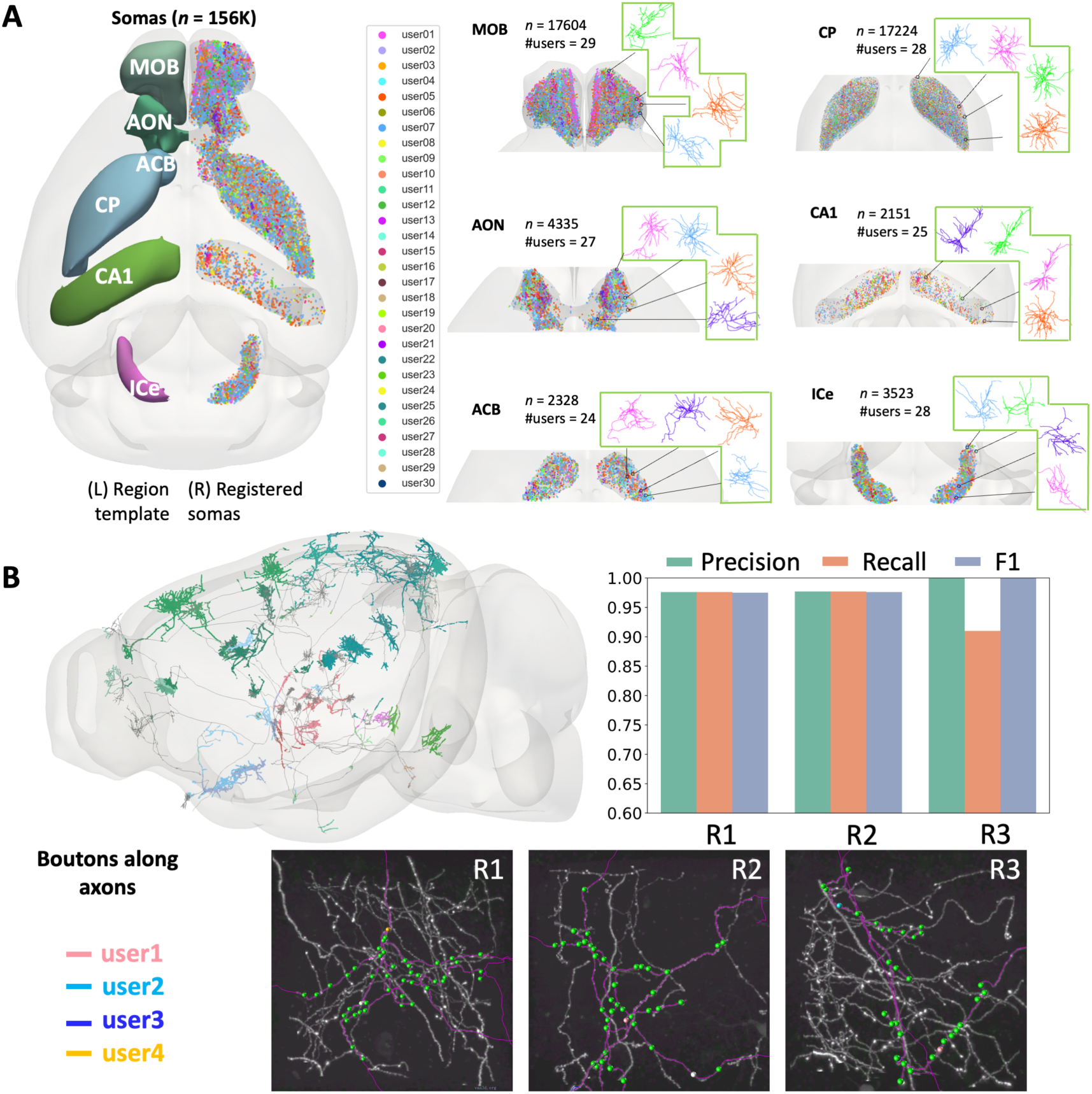
Reconstruction of sub-neuronal structures using CAR. **A,** The left panel: On the right side, a top-down view (CCFv3) of 156k semi-automatically annotated somas is presented, depicting eight selected brain regions color-coded along the anterior-posterior (AP) axis (left). Corresponding soma locations are plotted as dots (right). The different colors of the somas represent different users who have contributed to the annotations. The right panel: This panel provides close-up views of the eight selected regions mentioned in the left panel. Auto-traced dendrites are also shown for those neurons whose somas have been annotated by different users. The eight selected regions include: MOB (Main olfactory bulb), AON (Anterior olfactory nucleus), SI (Substantia innominata), CP (Caudoputamen), CA1 (Field CA1), ICe (Inferior colliculus, external nucleus), CUL4,5 (Lobules IV-V), SIM (Simple lobule). The panel displays the numbers of annotated somas and users involved in the region. **B,** The top-left panel showcases the sagittal view of 20 neurons with boutons that have been generated and proofread by CAR. Each distinct color represents boutons from different brain regions. The bottom panel consists of three image blocks (maximum intensity projection onto 2D is showed), denoted as R1, R2, R3, which have been selected for evaluation. Within the panel, potential presynaptic sites that have been auto-detected are marked with green markers, while white markers are identified as incorrect boutons. Boutons validated by 4 users are depicted using colors other than green or white, highlighting their agreement in the annotation process. In the top-right panel, the precision, recall, and F1 scores for these three selected image regions validation results are presented. These scores reflect the accuracy and completeness of the synapse validation results made by CAR users.

A striking advantage of utilizing the CAR platform lies in its ability to streamline complex brain image analysis, spanning from the whole-brain scale down to the level of synapses connecting neurons. This advantage is exemplified in the case of whole-brain axon tracing (**Fig. 2**), followed by the detection of axonal boutons which are the potential presynaptic sites of neuron connections (**Fig. 5B**). These boutons frequently manifest as concentrated varicosities arranged along axonal tracts, exhibiting an uneven distribution pattern (Liu, et al., 2023; Qian, et al., 2023). Detecting or validating boutons directly, without any spatial constraints, would pose a formidable challenge (Peng, et al., 2011). However, CAR’s precise axon tracing and reconstruction of spherical image objects, such as somas, greatly alleviate the challenge of bouton validation. The guidance provided by neurite fibers lends valuable cues for confirming boutons, and these structures can also be visualized using CAR’s toolkit. Consequently, we successfully examined brain-wide bouton distributions in conjunction with fully reconstructed neurons (**Fig. 5B**). We randomly selected three image regions, each sized at 256×256×256 pixels (117.76×117.76×512 μm), designated as R1, R2, and R3, each containing both axon tracts and numerous boutons that were verified together by four individuals using CAR. Both the precision and F1 scores (**Methods**) exceeded 0.9, affirming the suitability of CAR for comprehensive, large-scale analytics of whole-brain morphometry.

## Discussion

While recent endeavors have showcased achievements in acquiring thousands of complete mouse neuron morphologies (Peng, et al., 2021; Winnubst, et al., 2019) and developing several valuable software tools in the process (Ai-Awami, et al., 2016; Economo, et al., 2016; Peng, et al., 2014a; Peng, et al., 2014b; Wang, et al., 2019), the task of generating high-quality morphometry on a large scale remains a challenge yet to be fully resolved. Neuron reconstruction and other morphometry acquisition problems present several significant hurdles. Within these challenges, establishing the accuracy of neuron morphology is a complex endeavor, owing to the inherent intricacies of neurons and the potential impact of individual annotator biases. The pursuit of heightened accuracy often raises concerns about efficiency. Within our study, we confront this challenge by introducing CAR, a pioneering neuroscientific tool designed to foster immersive collaboration and facilitate the rectification of morphological and topological errors. CAR takes a distinct approach by harnessing the potential of *collaborative human intelligence augmented by artificial intelligence*. Through the integration of expertise and insights from multiple participants, our tool strives to achieve reconstructions that not only align with biological realities but also garner consensus among collaborators. We acknowledge the pivotal significance of faithfully capturing the intricate intricacies of neural morphology, all while ensuring a representation that genuinely reflects the underlying biology.

A central advantage of CAR lies in its capacity to facilitate thorough evaluation and validation of reconstructed data. Through its provision of immersive interaction and collaborative editing of neuron anatomy, CAR empowers researchers to seamlessly collaborate, capitalizing on their combined knowledge and expertise. The platform offers both accessibility and flexibility, enabling researchers situated in different locations or laboratories to collaborate in real-time. By merging annotations and contributions, participants can heighten the accuracy and comprehensiveness of reconstructions, surmounting the constraints of individual endeavors. Multi-collaboration not only elevates the quality of reconstructions but also fosters the exchange of knowledge and accelerates advancements in the realm of neuronal morphology analysis.

Queries regarding the efficacy of a multi-party collaboration within a multi-dimensional space to enhance tasks are deserving of further investigation. FlyWire (Dorkenwald, et al., 2022) endeavored to collaboratively proofread neural circuits using a browser-based interface with spatially chunked supervoxel graphs. However, the performance of the browser-based interface could present potential challenges and limited scalability when handling extensive datasets. In contrast, the CAR framework incorporates a range of heterogeneous devices, including personal computers, VR headsets, and mobile phones, each offering distinct advantages tailored to specific tasks, all supported by the CAR cloud server. Mobile clients are particularly well-suited for lightweight scientific tasks, offering convenient data visualization and sharing capabilities. VR platforms, on the other hand, excel in tackling intricate neuron annotation tasks, such as reconstructing neurons characterized by varying image qualities and densely clustered structures in noisy images. The inclusion of a game console adds an interactive, gamified element that engages users and motivates increased involvement in the reconstruction process. The compatibility of these diverse devices facilitates efficient visualization and annotation of complex 3D neuroscience data. By providing these varied client options, the framework leverages the strengths of different devices to contribute to data validation, substantiating its completeness and accuracy.

Efficiency stands out as another notable advantage of CAR. To optimize efficiency, CAR integrates AI tools like the branching point verifier and breakpoint detector. These AI agents collaborate with users, providing continuous assessment of user-generated reconstructions and furnishing reconstruction suggestions directly within the CAR platform. By harnessing computational capabilities, these AI agents amplify human contributions, elevating the precision and efficiency of reconstructions. The fusion of AI automation components and multi-collaboration within CAR empowers annotators to visualize and manipulate 3D neural reconstructions more effectively. This streamlined workflow significantly reduces the time and effort required for precise annotation without compromising the biological authenticity of the reconstructed morphologies. Furthermore, owing to the heterogeneous nature of CAR clients, AI can also be employed for AI-based data fetching, data I/O and transportation, data management across different clients, as well as innovative data generation, sharing, and entertainment. By harnessing the potential of AI and fostering collaborative endeavors, CAR not only enhances the efficiency of reconstruction processes but also elevates the overall user experience across a spectrum of data handling and interaction facets.

Thanks to the distinctive attributes of CAR outlined earlier, it serves a diverse array of neuroscience applications. Particularly, we harnessed CAR to explore brain anatomy across multiple scales, engaging in tasks such as annotating 3D brain regions, reconstructing neurons spanning entire mouse or human brains, delineating local dendritic and axonal arbors, pinpointing somas, validating potential synaptic sites, and performing a variety of morphometric measurements. The accomplishments achieved through the CAR platform in these applications highlight its potential to revolutionize future undertakings focused on cell types and brain anatomy. This transformative potential contributes to the advancement of our comprehension of the intricate neuronal networks that underpin a multitude of biological processes.

Notably, Woolley et al., (2010) presents empirical evidence highlighting the emergence of a collective intelligence factor in group collaboration. The study underscores that a group’s collective intelligence isn’t solely tethered to the individual intelligence of its members. Instead, elements such as social sensitivity, equitable participation, and gender diversity exert a more substantial influence on collective intelligence. These findings carry significant implications for comprehending group dynamics and efficacy. They underscore that effective group dynamics and interactions are pivotal for fruitful collective decision-making and problem-solving. When we developed CAR, we noted that drawing a comparison between crowd wisdom and individual decision-making could yield several key insights. While individual decision-making can be susceptible to biases and a limited perspective, crowd wisdom amalgamates diverse viewpoints, mitigating individual biases and offering a more encompassing perspective conducive to accurate judgments and solutions. Crowd wisdom also capitalizes on the diversity inherent in a group, encompassing a broader array of experiences, knowledge, and skills. However, we also note that crowd wisdom doesn’t guarantee superior outcomes across all scenarios. Factors like groupthink, undue reliance on popular opinion, lack of diversity, and suboptimal group dynamics can undermine its efficacy. Hence, cultivating an environment that nurtures diverse thinking, balanced participation, and positive social dynamics becomes imperative for successful engagement with crowd wisdom. These latter situations should be paid attention to when we apply CAR in real applications.

Looking into the future, we envision broader applications for CAR in various biological contexts, granting users the capability to immerse themselves in multi-dimensional volume exploration environments while benefiting from an array of AI tools. The aptitude of CAR to perform neuron morphometry for Big Data in brain science has unlocked promising avenues for essential research endeavors. These encompass intricate cell typing paradigms (Munoz-Castaneda, et al., 2021; Peng, et al., 2021) and the potential establishment of connectomes through the utilization of light-microscopic brain images (Sporns, et al., 2014; Xiong, et al., unpublished private communication).

## Acknowledgement

This work was mainly supported by a Southeast University (SEU) initiative of neuroscience awarded to H.P.. The Southeast University team was also supported by a STI2030-Major Projects Grant No. 2022ZD0205200/2022ZD0205204 awarded to L.L.. Y.W. was supported by the National Natural Science Foundation of China (32071367), the Guangdong High Level Innovation Research Institute (2021B0909050004), and the Key-Area Research and Development Program of Guangdong Province (2021B0909060002).

## Author contributions

H.P. conceptualized and managed this study, and instructed the detailed development of experiments. Y.W., L.H., L.Z., Y.Z., and Y.H. developed the CAR client and server software. Y.W. conducted the experiments with help of L.Z., Z.Y. coordinated the data annotation, and L.Z., Z.Y., Y.Z., Y.H., K.L., and L.W. analyzed the data. L.Z., Z.Y., Y.Z., J.X., Z.G., D.C., L.Le, J.C., H.Zeng, W.Y. and H.Zhang participated in the experiments. L.Liu and X.C. contributed the preparation of imaging datasets of mouse and human, and also provided assistance in data curation. H.P., L.Z., and Y.W. wrote the manuscript with assistance of all authors, who reviewed and revised the manuscript.

## Data availability

The mouse and human neuron reconstructions released are deposited to https://github.com/neurogeom/CAR/tree/main/data.

## Code availability

CAR is available at GitHub, https://github.com/neurogeom/CAR, with both source code and binary executables.

## Methods

### Mouse brain regions abbreviation

**Isocortex:** primary motor area (MOp), secondary motor area (MOs), primary somatosensory area, mouth (SSp-m), supplemental somatosensory area (SSs), ventral auditory area (AUDv), primary visual area (VISp), ventral part (RSPv).

**Thalamus (TH)**: ventral posterolateral nucleus (VPL), ventral posteromedial nucleus (VPM), anteromedial nucleus (AM), reticular nucleus (RT), lateral dorsal nucleus (LD), lateral geniculate complex, dorsal part (LGd), paraventricular nucleus (PVT), medial geniculate complex (MG), lateral posterior nucleus (LP), lateral dorsal nucleus (LD).

**Cerebral nuclei (CNU):** caudoputamen (CP), olfactory tubercle (OT), substantia innominata (SI).

### Human brain regions abbreviation

Frontal lobe (FL); superior frontal gyrus (SFG); middle frontal gyrus (MFG); inferior frontal gyrus (IFG); superior temporal gyrus (STG); middle temporal gyrus (MTG); parietal lobe (PL); inferior parietal lobule (IPL); temporal pole (TP); occipital lobe (OL); an intermediate region bordering superior frontal gyrus, middle frontal gyrus and inferior fronton gyrus (S(M,I)FG).

### Collaborative neuron reconstruction protocol

To facilitate flexible and organized collaboration among CAR users, we have devised a straightforward yet highly effective neuron reconstruction protocol (see **Supplementary Fig. 3**). The protocol is underpinned by a set of rules governing the reconstruction process:

1. A user is permitted to annotate a neurite if it originates from one of the following: a) The soma. b) Another neurite previously reconstructed by the same user. c) A neurite that has been validated and confirmed.
2. Alternatively, users have the option to confirm, delete, or modify neurites previously reconstructed by other users, provided that these neurites either originate from the soma or extend from another already confirmed neurite.
3. It is essential to note that a user is precluded from confirming their own reconstructions, emphasizing the importance of impartial validation.
4. The neuron reconstruction process is considered complete only when all reconstructed neurites have been duly confirmed, and there are no further unaccounted structures that can be added.

By adhering to this protocol, we establish a robust framework for collaborative neuron reconstruction and verification. Annotations made by one annotator can be rigorously reviewed and endorsed by another annotator, thus bolstering the accuracy and reliability of the overall annotation results.

### Real-time collaboration design

We propose a real-time collaborative annotation system for neuron reconstruction, which operates on a cloud-based server. The collaboration process begins with the creation of collaboration rooms, where each room corresponds to a specific reconstruction task. These rooms are assigned unique identifiers and can be synchronized across multiple devices, including VR, desktop, and mobile platforms. The server, hosted on a public cloud, establishes a TCP Socket connection with all connected clients. Its main role is to receive and process annotation operation instructions from clients and broadcast them to other clients within the same collaboration room. This ensures that everyone has access to the same information in real-time. The CAR server timestamps these operations to enable sorting based on time on the client side. To enable simultaneous co-reconstruction by multiple users, the system implements a lock mechanism to prevent conflicts. Additionally, the system supports undo-redo functionality, allowing clients to revert previous operations and restore previous states.

### Analysis on mouse neuron reconstructions

1. Time stages split

To obtain temporal data, we utilize the log records of the CAR system, which enable the recovery of neuronal reconstruction results at any given moment. In order to analyze the structural patterns of the 20 neurons at a temporal resolution, we divide each neuron’s total reconstructed length into eight equal segments, resulting in eight time intervals. This approach allows us to analyze different neurons within the same temporal scale.

2. Normalized topological height (NTH)

Before calculating the normalized topological height, we ensure precise alignment of all axonal neurons using mBrainAligner for registration to CCFv3.

(1) To derive the topological height, we consider the terminal nodes as having a height of 1. As we traverse the neuron structure, the height of each branching point is determined by adding 1 to the highest level among its child nodes (**Supplementary Fig. 7**).
(2) In Step 2, we expand on this process by setting the maximum level observed among eight time points as the denominator for calculating the topological height. Each topological height is then normalized as following:

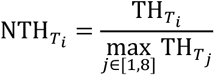 where a smaller level indicates a more peripheral position. This approach enables the comparison of forms across different time points.
(3) Furthermore, under each normalized topological height, we calculate the average length of both the matched and unmatched parts of 20 neurons at each time point.

3. Computation of consistency of axonal neuron

The consistency of neurons in Figure 2D is defined as the ratio of reconstructed morphology lengths that have been validated by other individuals (users) to the total length of the neuron morphology across all eight stages.

4. Computation of contrast projection map

The contrast projection map comprises two distinct components: the neurites that have been added through collaboration, denoted by the symbol ‘+’, and the neurites that have been subtracted through collaboration, represented by the symbol ‘-’. The subtract projection map represents the reconstructed neuron morphology that has been deleted by others.

Regarding the addition projection map, we conducted a comparison for each annotator at every branching point. If the annotator assigned to a leaf node within a branch differed from the parent nodes’ annotation, it indicated the addition of a neuron through collaboration.

### AI-based automation tools

Two AI-based tools in the user annotation process to assist users achieve complete neuron reconstruction by identifying feature points including the branching and the breakpoints of neurons.

1. Implementation of BPV and BD

To verify branching points, we designed a Convolutional Neural Network called RSHN (Residual Single Head Network). The network consists of a down-sampling module, an attention module, and two residual blocks. The network is designed to accept an image patch cropped from the whole brain image that is centered around the branching point as its input. To reduce the dimensionality of the input, the patch undergoes a down-sampling process. The down-sampling operation is achieved by applying two 5×5×5 convolution kernels with a stride of 1, followed by two 3×3×3 convolution kernels with a stride of 2. Following that, the network applies an attention module and residual blocks to extract significant features from the image patch. The residual block consists of two convolutional layers and one batch normalization layer. ReLU is used as the activation function for non-linear processing. Finally, the output is obtained through a fully connected layer for classification (**Supplementary Fig. 9A**).

To detect breakpoints, RDHN (Residual Double Head Network) networks, a variant of RSHN, are designed to process two inputs: an image patch extracted around the breakpoints and a corresponding mask image. The two images are separately down-sampled, and the resulting features are fed into the attention module for feature enhancement. The purpose of this is to emphasize the disparities between breakpoints and branching points by obscuring the shared areas. This approach guarantees that the network acquires more distinguishing features and enhances its ability to differentiate between the two types of points (**Supplementary Fig. 9B**).

2. Evaluation of BPV and BD

To evaluate the accuracy of the detection in the module, we designed below metrics for assessment. In this context, the final expert-proofed reconstruction is designated as the ground truth (GT). For candidate branching points, they are selected from the current reconstruction. These points serve as the central reference to extract image patches, which are subsequently used as inputs for the classifier. Similarly, for candidate breakpoints, they are chosen from all the terminations found in the current reconstruction. The image patches surrounding these points are also extracted and utilized as inputs to the classifier, with the corresponding breakpoints serving as the centers.

|  | BPV | BD |
| --- | --- | --- |
| **TP** | The number of branching points that exist in the GT and are classified as branching points | The number of breakpoints that are missing in the GT and hinted as breakpoints |
| **TN** | The number of branching points that are missing in the GT and hinted as non-branching points | The number of breakpoints that exist in the GT and are classified as terminations |
| **FP** | The number of branching points that are missing in the GT but classified as branching points | The number of breakpoints that exist in the GT but are hinted as breakpoints |
| **FN** | The number of branching points that exist in the GT but hinted as non-branching points | The number of breakpoints that are missing in the GT but classified as terminations |

The accuracy (ACC) of the classification model is calculated as follows:

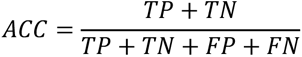

### Analysis on human neuron reconstructions

Before analyzing human neuron reconstructions, all neurons underwent preprocessing steps. All neuron morphology has been preprocessed by implementing sorting, rescaling to 1 pixel to 1μm space, and resampling to ensure reasonable analysis results.

1. User attentions

The computation of user attentions involves several steps. Initially, a cuboid region with dimensions of 20×20×20 μm³ is defined as a bounding box surrounding each neuron node. Within these regions, we collect the unique user editing that occur. To analyze user attentions, we consider the number of distinct users identified and assign this count as a ‘type’ attribute to each node. The number of ‘types’ corresponds to the users who contributed to either adding or modifying the neuron segment.

2. Local structural complexity

Our method for computing local structural complexity builds upon the concept of Sholl analysis but expands its application by considering not only the soma but also each individual point as a center for calculating intersections with surrounding points.

3. Signal complexity

To compute signal complexity, we utilize the reconstructed morphology of neuron and estimated radius values as masks. Each voxel in the volume image is classified as either foreground or background based on these masks. Subsequently, the image is decomposed into a number of small cubes, e.g., 20×20×20 in size. For each cube, its signal complexity is defined as the mean value of the foreground voxel intensity divided by the mean value of the background voxel intensity. Additionally, by uniformly converting the signal complexity values into the range of [0,255], we can generate a specialized 3D image that visually represents the signal complexity of the original image.

### Analysis on human neuron image

To assess the image quality of human neuron images in Supplementary Figure 11A, an image decomposition method called non-negative matrix factorization (NMF) is utilized (Févotte and Idier, 2011). In the NMF decomposition process, the average of every 10 image slices along the z-axis is calculated and transformed into 1-dimensional vectors. These vectors are collected into a matrix, on which a 3-component NMF model is constructed to obtain the decomposed components representing the background and the signal obtained as the difference between the image block and the background component, enabling better separation between the two. After performing the NMF decomposition, several metrics are calculated: ‘signal.median’: This metric refers to the median value of the signal. ‘signal.rsd’ (relative standard deviation): It is calculated by dividing the standard deviation of the signal by the median value of the signal. ‘contrast’: This metric is obtained by subtracting the median value of the signal from the median value of the background. It is important to note that during the calculation, only the background values below the 99th percentile and the signal values above the 90th percentile are considered, which helps to reduce the impact of residual signal in the estimated background and vice versa.

### Soma-pinpointing in CAR-Mobile

The soma identification protocol in CAR involves two major steps. The first step is the automatic detection of potential soma positions on the CAR server. In this step, by applying the gray-scale distance transform (GSDT) algorithm to the brain image, we identify voxels with intensities ranging from 5 to 30. These voxels are considered as potential soma candidate positions.

The second step involves pinpointing somas in CAR-Mobile client. The process goes as follows:

1. When a user wants to open a file, they simply click the “open file” button in the mobile client.
2. The client sends a message to the server requesting potential location information.
3. The server checks the potential location table and selects an unprocessed location record for the client.
4. The client then sends a message to the server, requesting an image block centered around the location information with the appropriate size.
5. Additionally, the client sends another message to obtain existing soma positions within the bounding box of this specific block. It’s important to note that image blocks may overlap when potential locations are close together, so somas uploaded by other users may appear in the client’s block.
6. The server crops the image block from the whole-brain image based on the requested location and size.
7. The server also looks up the existing somas relevant to the client’s request. Clients are empowered to update soma information by making changes, additions, or corrections to the identified soma data.

CAR-Mobile incorporates optimization techniques to facilitate efficient online collaboration. To prevent conflicts arising from simultaneous access to the same image, the CAR server implements a locking and expiration strategy. When an image is distributed to a client, the corresponding record in the table is locked, preventing the image from being distributed to other clients while the lock is active. The lock is automatically released when the client returns the annotation result or after a predefined period of 8 minutes.

To enhance the browsing experience, CAR-Mobile utilized a pre-download strategy. It maintains a queue of images, and a dedicated download thread ensures that the queue remains populated. When a user requests an image, the first image in the queue is retrieved, and any newly downloaded images are appended to the end of the queue. Each downloaded image had a predefined expiration time of eight minutes from its initial download. Once expired, the client could no longer perform any actions with the image. This optimization strategy allows for efficient resource allocation and provides a smoother browsing experience within the CAR system.

### Synapse validation in CAR-Mobile

An axonal bouton, which is characterized as a localized swelling along axonal shafts, manifests as a region of high intensity in light microscopy data when observed at a submicron resolution. The synapse validation process bears similarities to the process of pinpointing somas in CAR-Mobile client. It also consists of two steps:

Firstly, we utilize an algorithm based on the approach presented in the publication by Liu et al. (2023) to automatically detect potential bouton positions. This method combines intensity and radius profiles along axonal shafts to identify initial candidates for boutons, characterized by overlapping peaks in both profiles. False positives are eliminated using heuristic criteria: boutons should be 1.5 times larger than surrounding nodes, have intensity values above 120 in 8-bit images, and duplicate candidates closer than 5 voxels are discarded.

Next, we crop image blocks sized at 128×128×128, and its corresponding candidate synapses as well as morphology results. These blocks along with synapses and morphology results are distributed to clients, where users engage in proofreading tasks to identify and correct any missing or erroneous synapse sites within the image block. The validation results are sent back to the server and shared with other users for cross-validation. Each image block is distributed to two individuals for this purpose.

Secondly, validation process in CAR-Mobile client. the pre-detection of boutons and the corresponding morphology results are integrated into CAR-Mobile clients for rendering. As a result, users only need to engage in proofreading tasks, identifying and correcting any missing or erroneous synapse sites within an image block distributed from the server. The validation results are then sent back to the server and distributed to other users for cross-validation. Each single image block is distributed to two individuals for this purpose.

## Supplementary Tables

**Supplementary Table 1.**
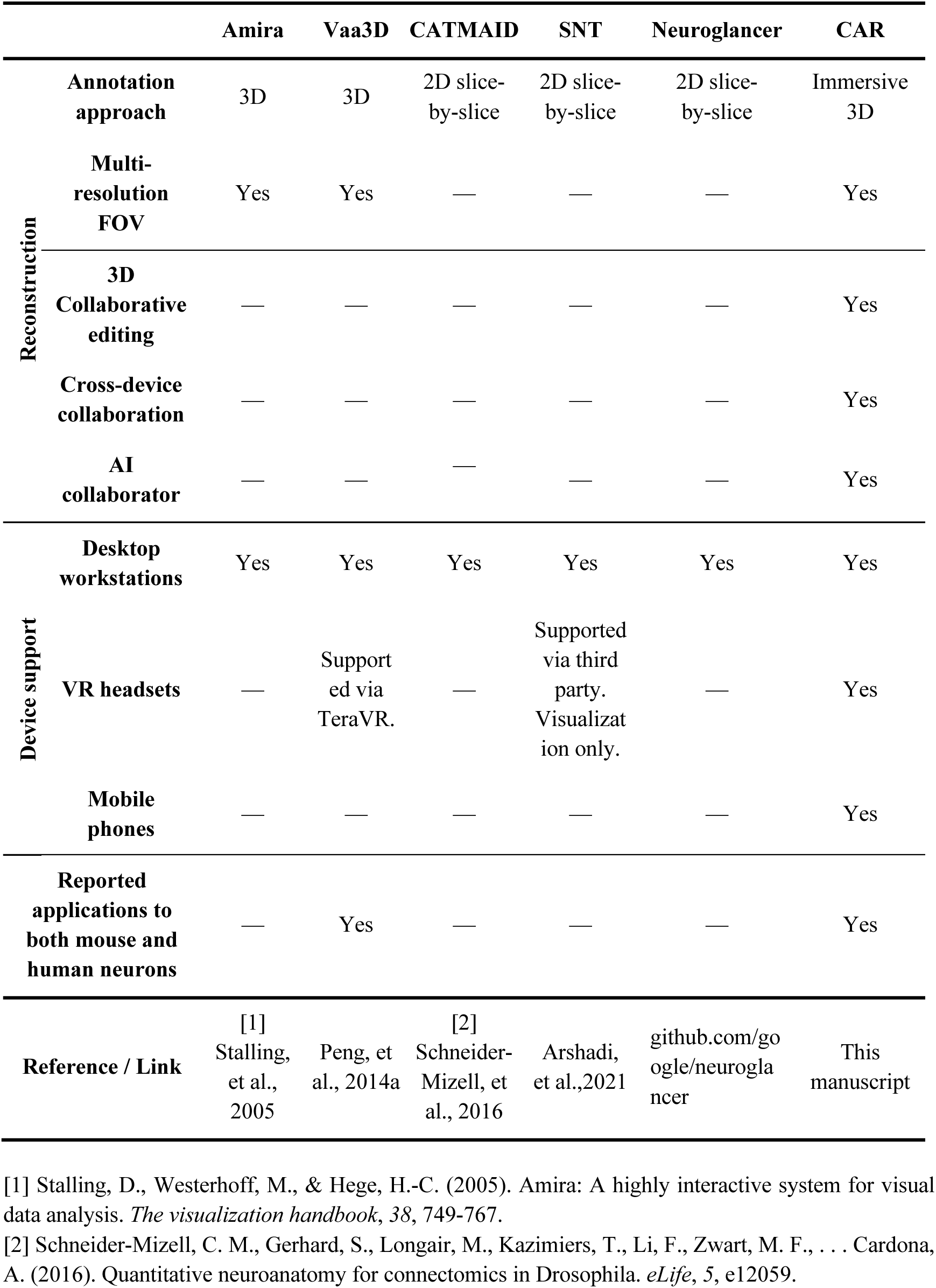
A comparison between CAR and other software tools for neuron reconstruction. In detail, we evaluated their capabilities on collaborative reconstruction and support for different kinds of devices. “—” indicates that the function is not supported by the specific tool.

**Supplementary Table 2.** Several exemplar CAR applications at various scales and the corresponding supported clients. “√” indicates the application is supported by the client, while “⛤” indicates the application is recommended to be performed on the client.

| Scale | CAR Applications | Clients |  |  |
| --- | --- | --- | --- | --- |
|  |  | CAR-WS | CAR-VR | CAR-Mobile |
| **1cm** | Brain region location | ☆ |  |  |
| **1mm** | Complete neurons/axons reconstruction | √ | ☆ | √ |
|  | Image quality evaluation and rating |  |  | ☆ |
| **100um** | Dendrites reconstruction | ☆ | ☆ | √ |
|  | Simple neurite structure proofreading | √ | √ | ☆ |
| **10um** | Soma pinpointing | √ | √ | ☆ |
| **1um** | Synaptic sites production/validation | √ | √ | ☆ |

## Supplementary Figures

**Supplementary Fig. 1.**
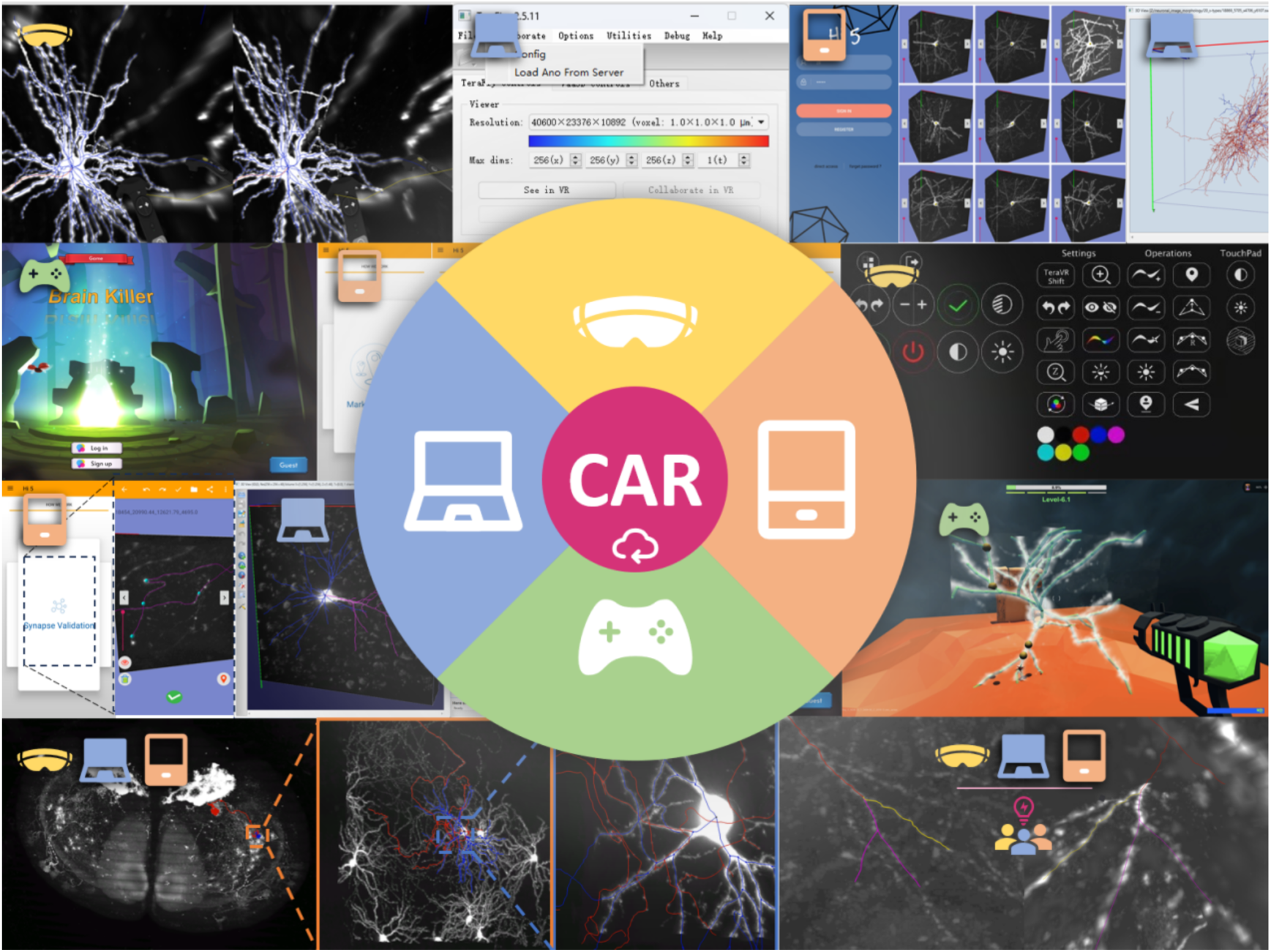
Overview of the CAR client software user interface. The CAR platform offers an array of diverse client software, each equipped with the ability to seamlessly interact with other clients, whether of the same type or different, through the CAR server. With the workstation (depicted by a blue icon), the virtual reality tool (yellow), or even the mobile app (orange), users can jointly explore neuronal data across multiple levels of resolution and execute various neuromorphometric tasks. Additionally, users may further access and curate neuronal data through the CAR game console (green).

**Supplementary Fig. 2.**
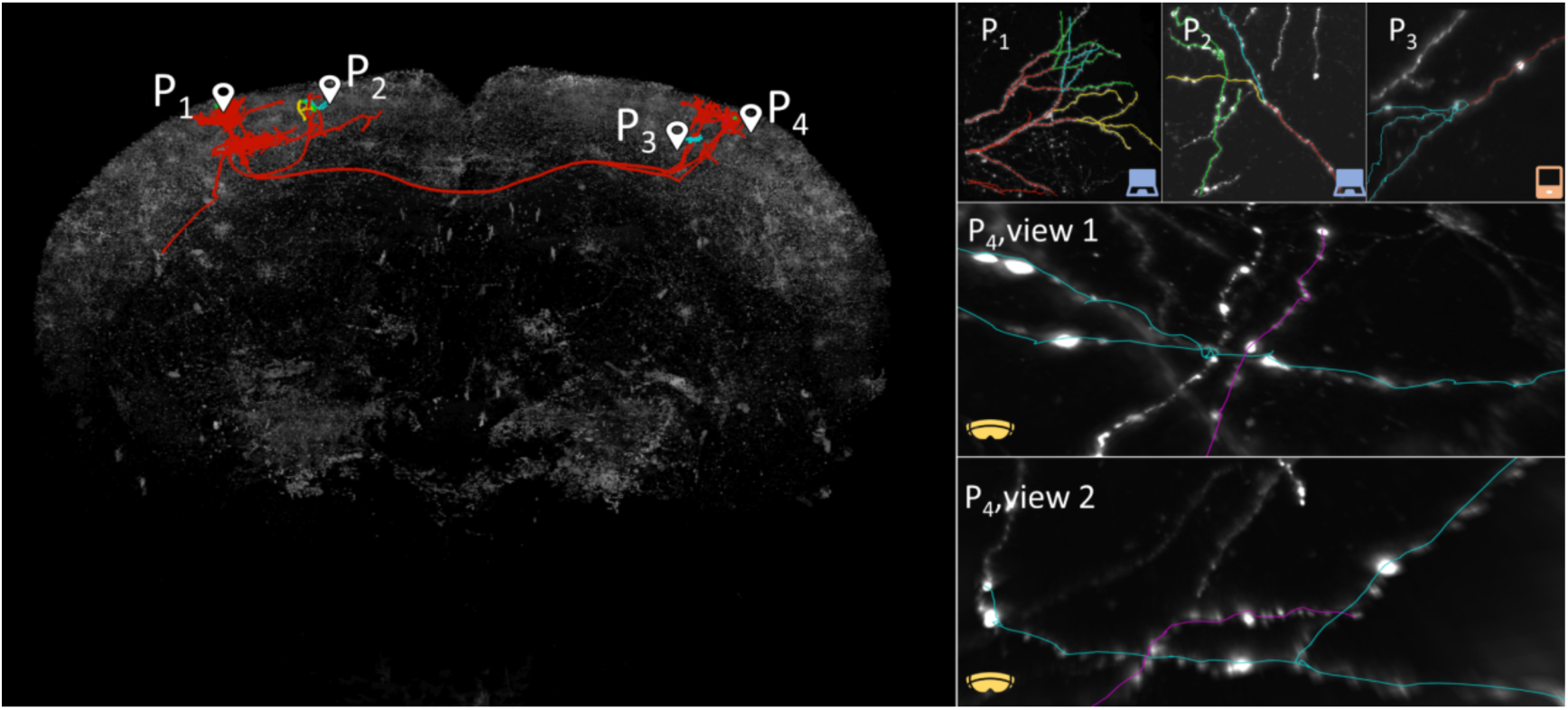
Exemplary collaborative neuron reconstruction from whole-brain data using CAR. In this exemplar collaborative effort, five users positioned at four locations (one user each at P1, P2, and P3, with two users at P4) utilized three types of CAR clients (desktop workstation, VR, and mobile app) to collectively reconstruct a neuron. The left panel provides a global view, while the right panel offers local perspectives. In all panels, neurites that have undergone proofreading are highlighted in red, while unchecked neurites are depicted in user-specific colors. Specifically, two users worked at P1 and P2, employing desktop workstations to reconstruct neurites. Meanwhile, the user at P3 inspected others’ reconstructions using the mobile app. At P4, two users wearing VR headsets collaborated to determine whether two adjacent neurites formed a bifurcation or not.

**Supplementary Fig. 3.**
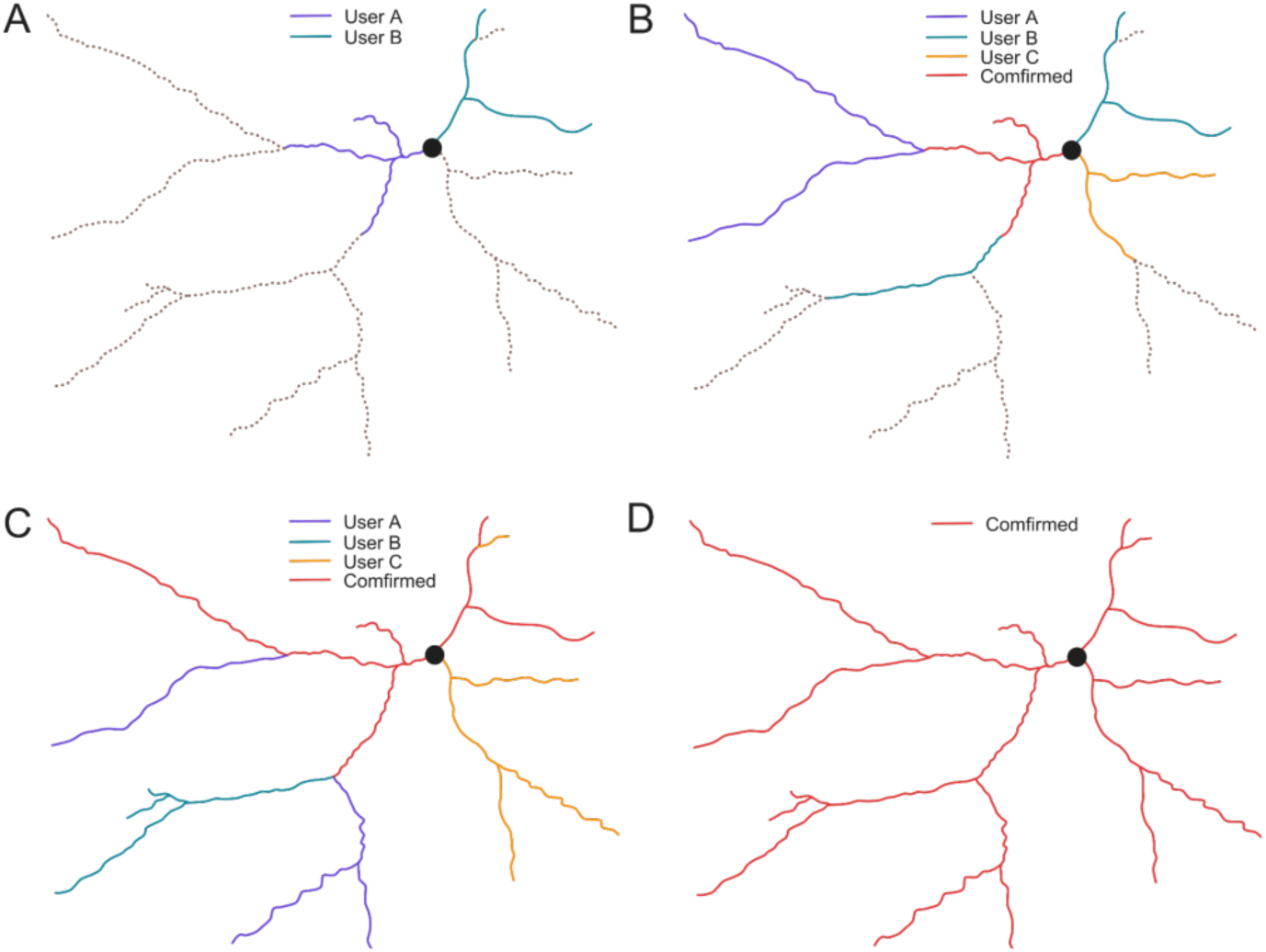
Collaborative neuron reconstruction protocol. **A**, **B**, **C**, **D**, four subsequent stages during the reconstruction of a neuron. Purple, green, and orange lines indicate neurites reconstructed by 3 different users, respectively. Red lines represent confirmed neurites. Dashed lines represent neurites that are yet reconstructed at the moment. A reconstruction is considered “finished” when all structures have been annotated and confirmed.

**Supplementary Fig. 4.**
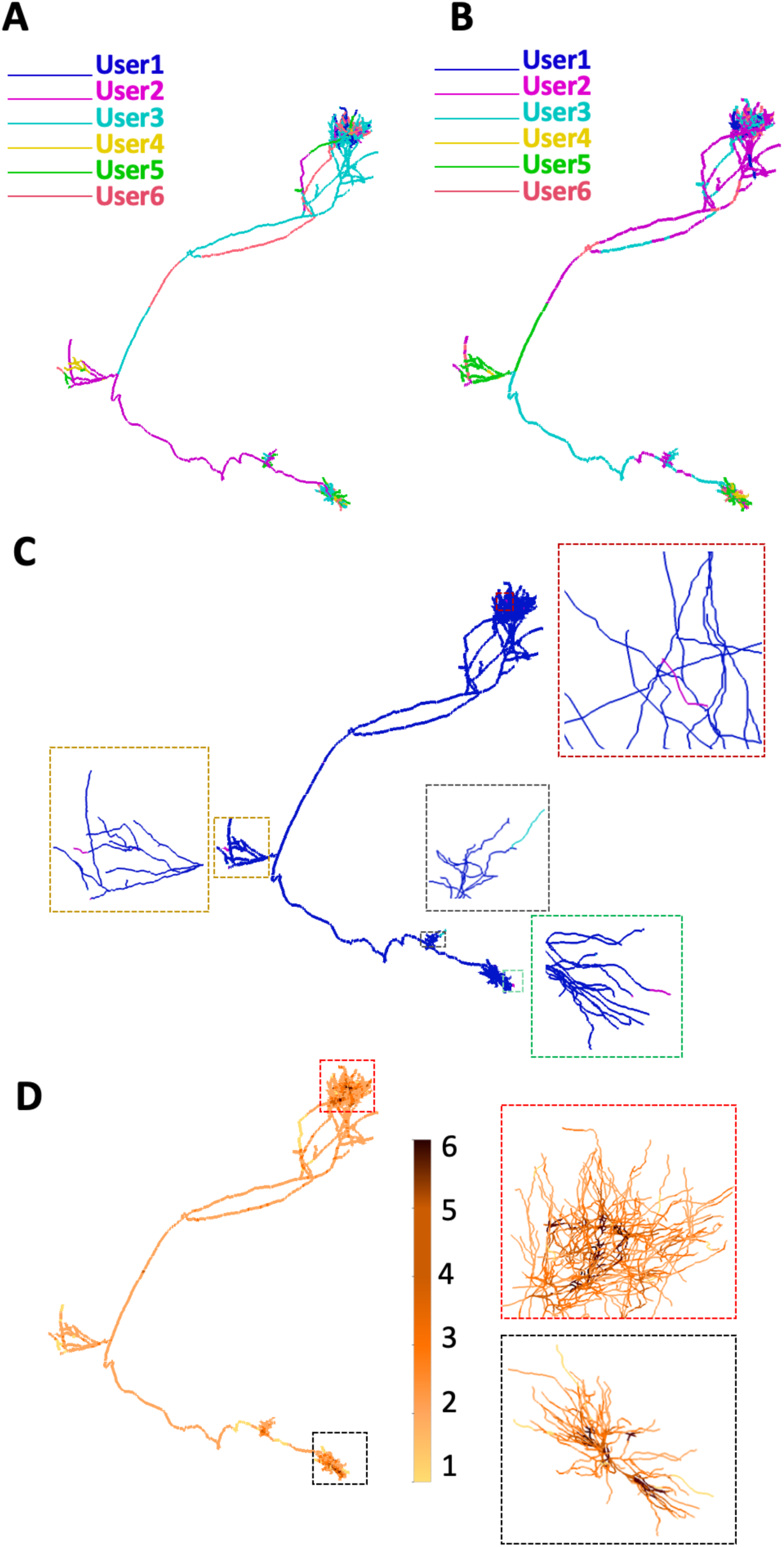
A CAR-reconstructed VPL neuron. **A**, Visualization of the neuron, with each user’s annotations represented in distinct colors. **B**, Visualization of the neuron, showcasing each user’s proofreading efforts in distinct colors. **C**, Expert modifications to the laymen’s consensus result. Neurites deleted by the expert are marked in magenta, while neurites added by the expert are highlighted in cyan. **D**, A heatmap illustrating user participation across various regions of the neuron. The number of participating users is color-coded according to the color map.

**Supplementary Fig. 5.**
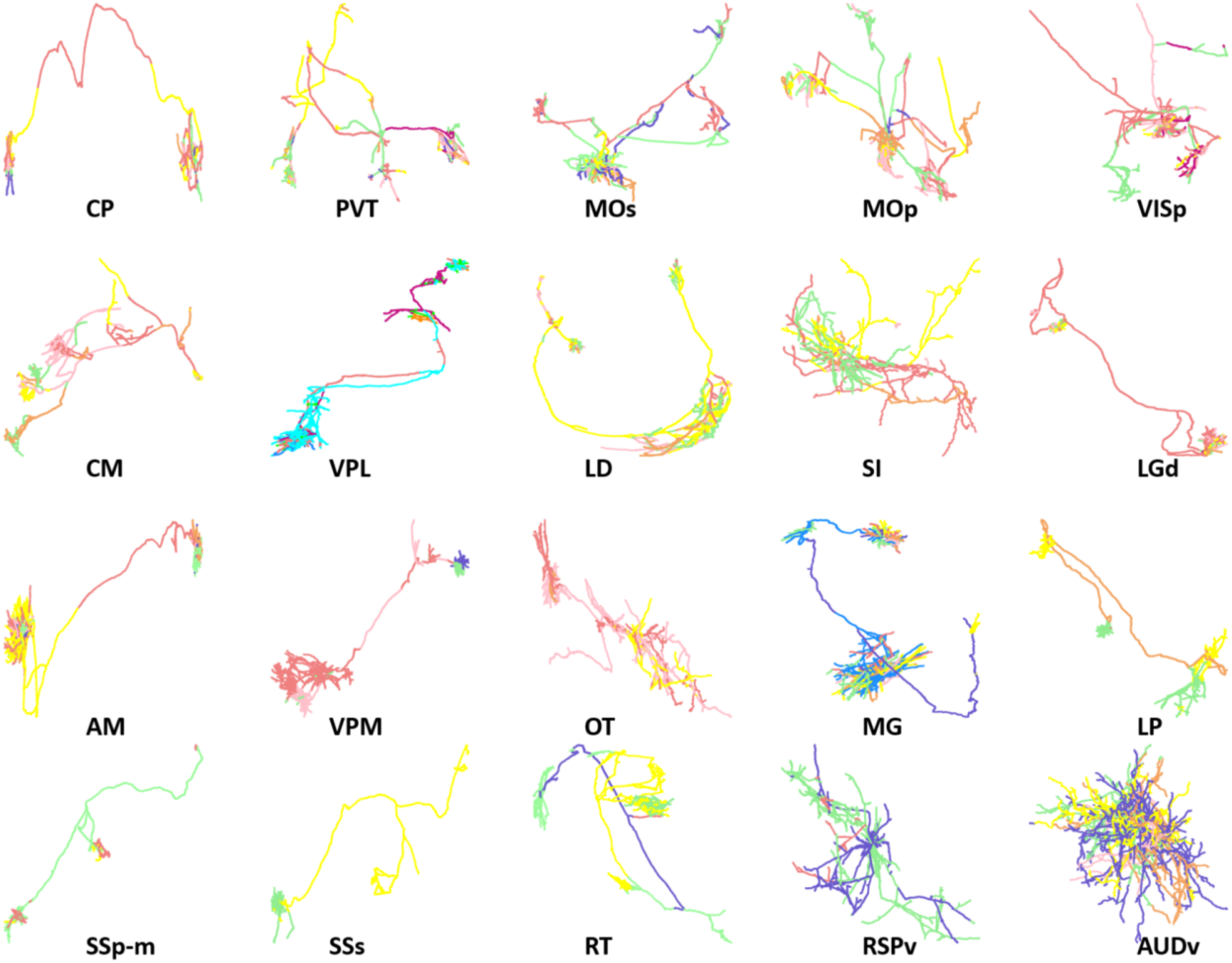
Collaborative reconstruction of 20 mouse neurons using CAR. In this collaborative effort, 12 individual users contributed to the reconstruction of 20 mouse neurons, while the number of contributors for each neuron varies. For an evaluation of their collaborative synergy, each user was assigned a unique color, and the neurons were color-coded accordingly to emphasize different users’ contributions. Additionally, the validation process may involve more users, who cross-check and verify neurite annotations made by others.

**Supplementary Fig. 6.**
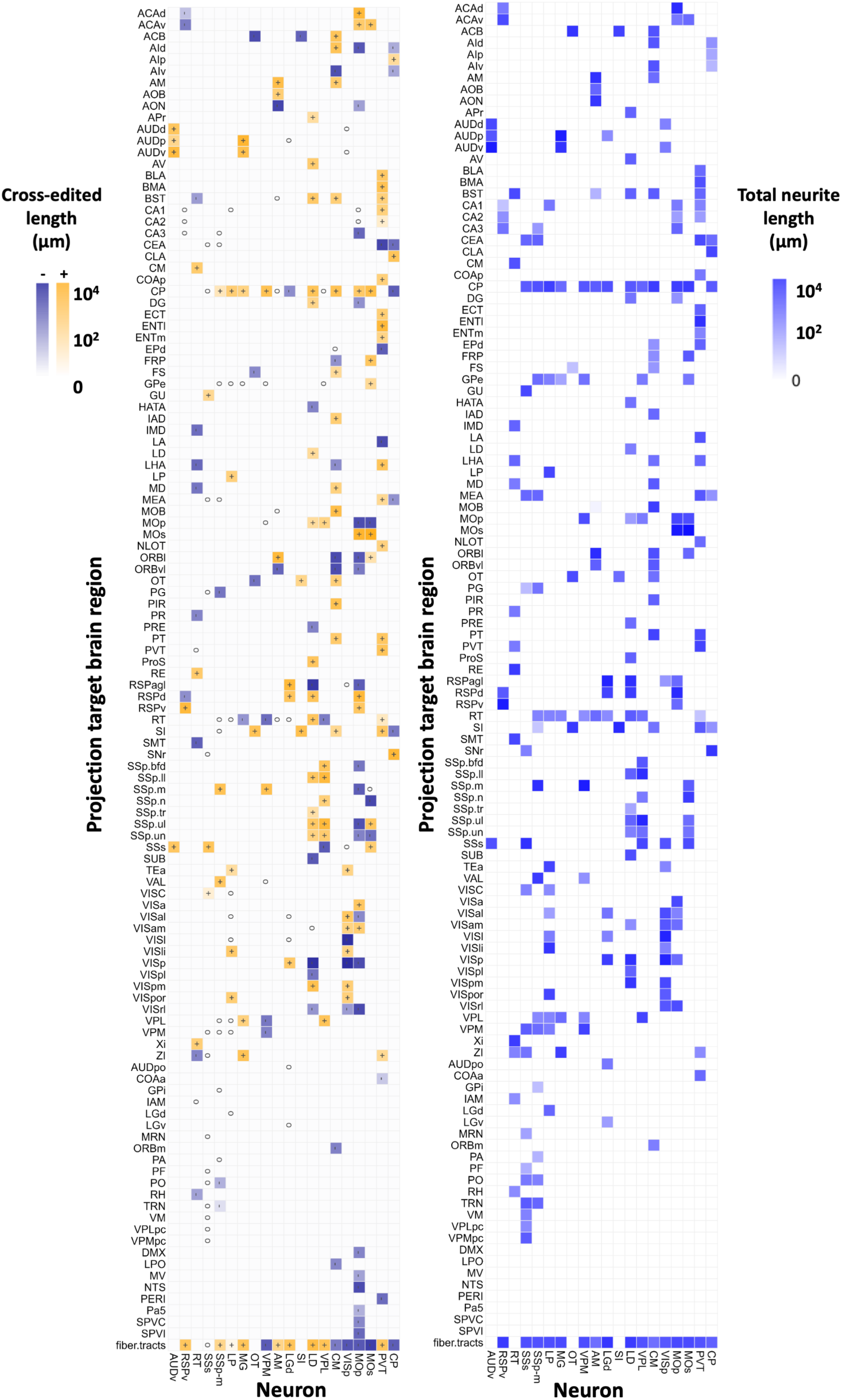
The projection maps formed by the 20 mouse neurons. **A**, A projection map derived from the collaboratively reconstructed sections of the 20 mouse neurons (identical to Fig. 2B, presented here again for comparison purpose). **B**, A complete projection map that encompass reconstructions from both the collaborative and non-collaborative efforts. Color coding reflects the total neurite length within each specific region.

**Supplementary Fig. 7.**
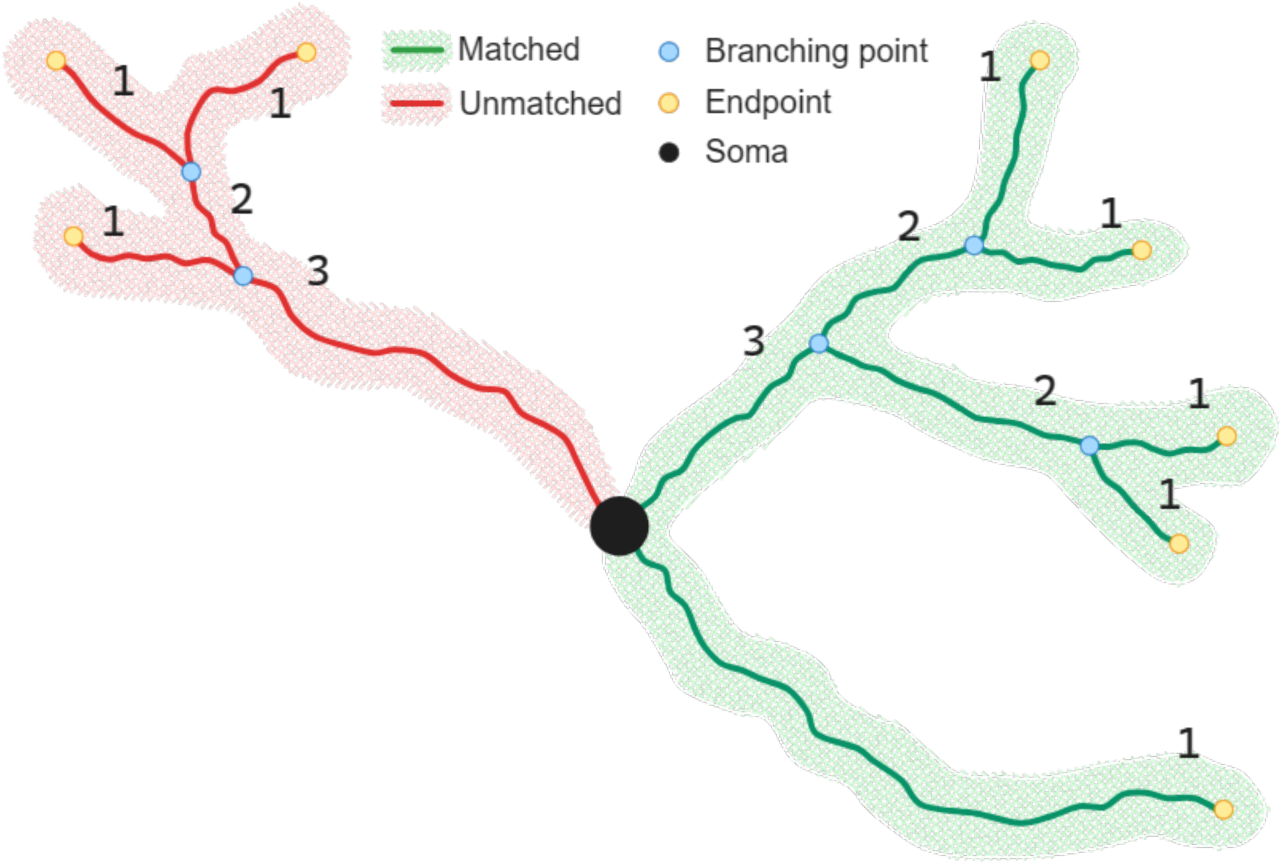
Illustration of the calculation of normalized topological height (NTH). Segments adjacent to endpoints are assigned a topological height of 1. For any other segment, its topological height is defined as one plus the greater topological height of its two sub-segments. Eventually, the topological height values are normalized by dividing them by the maximum topological height.

**Supplementary Fig. 8.**
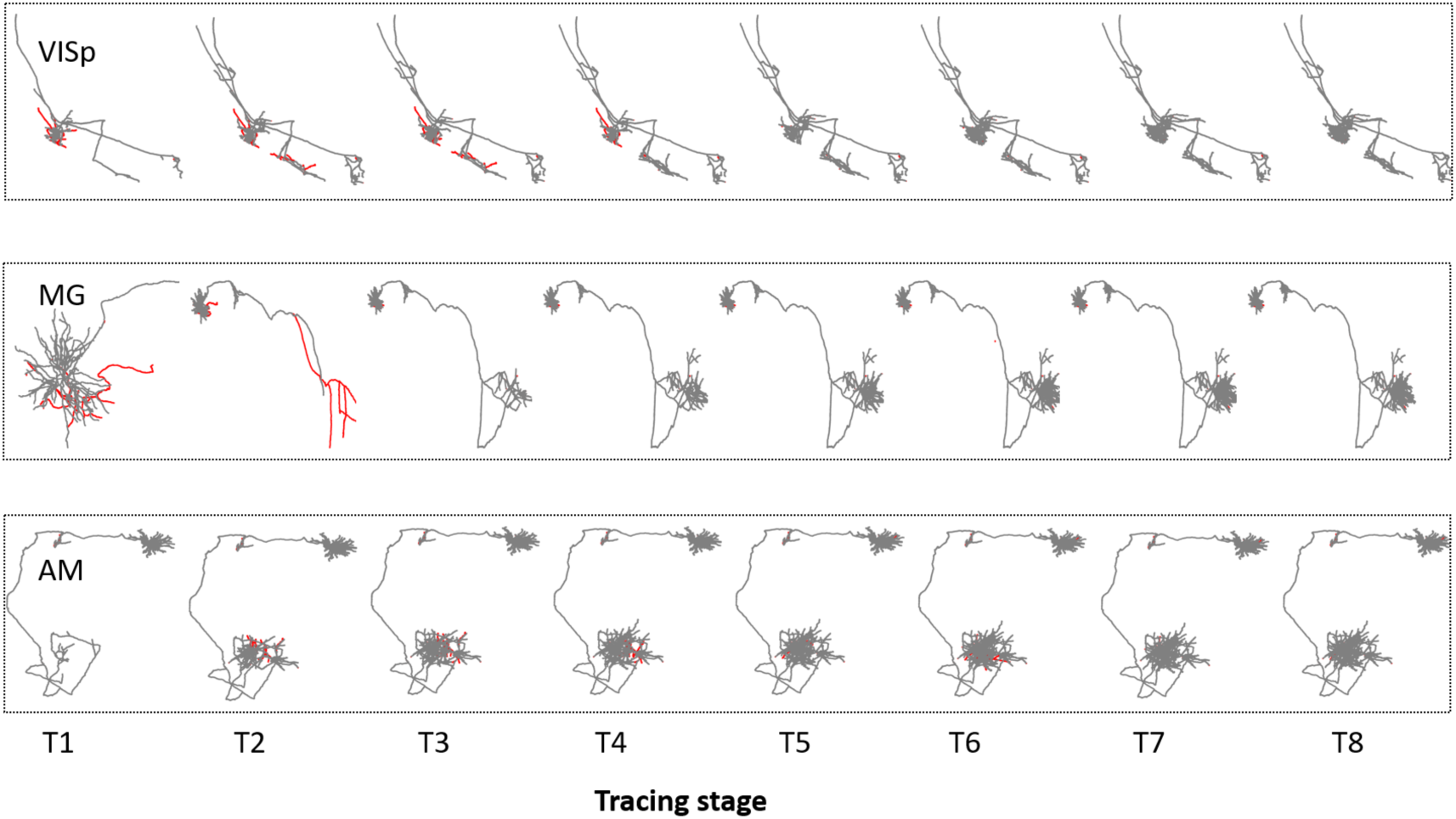
The morphologies across eight tracing stages for the reconstructions of VISp, MG, and AM neurons. Each row showcases a distinct neuron (VISp, MG, and AM), presenting its eight intermediate morphologies at time stages T1, T2, …, T8, arranged from left to right. Reconstructions in the early stages (e.g., T1, T2) may be scaled up for enhanced clarity. Neurites shown in grey color represent correct structures that are matched with the expert-validated reconstructions, while neurites shown in red color represent unmatched structures.

**Supplementary Fig. 9.**
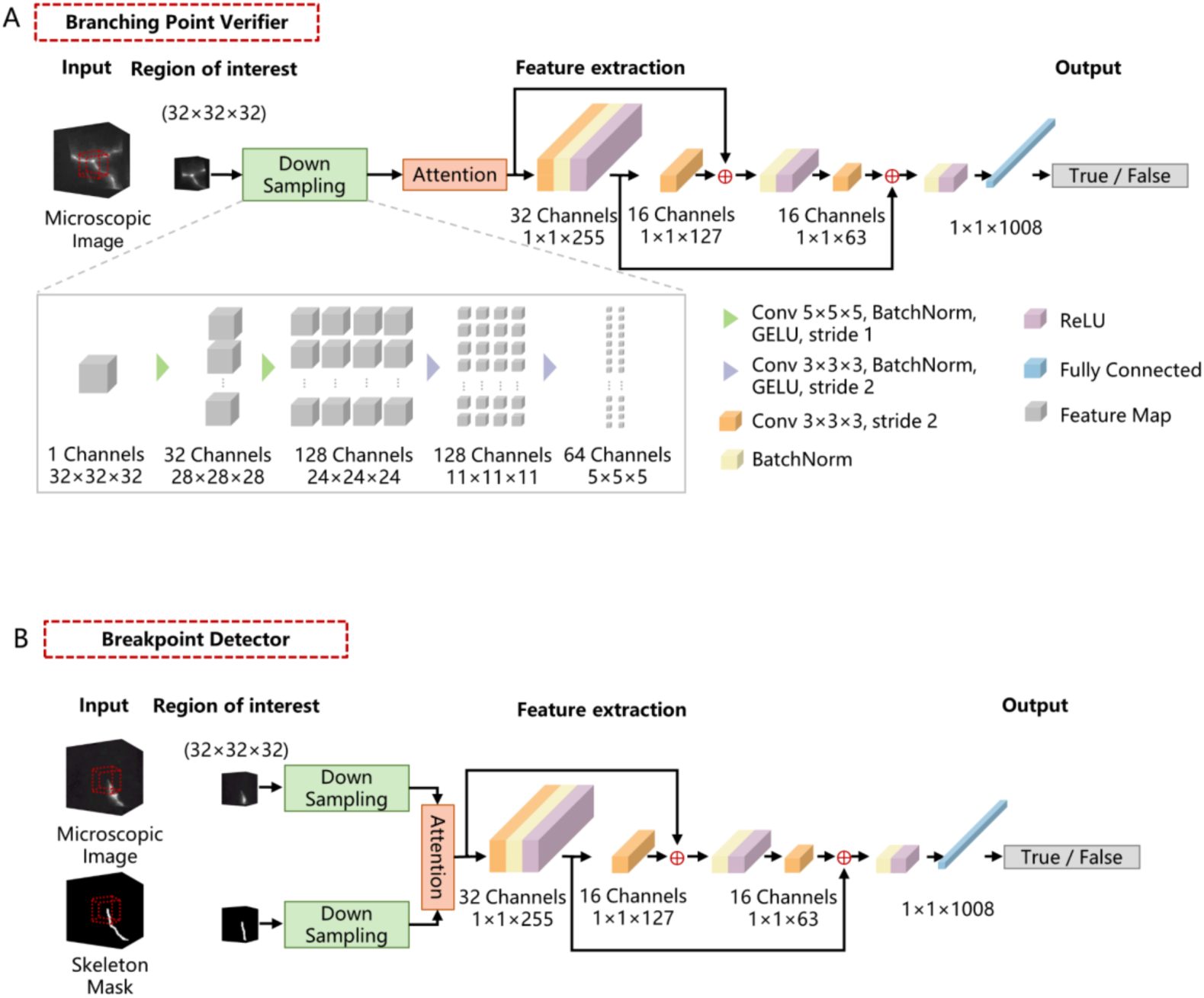
Network architectures of Branching Point Verifier and Breakpoint Detector. **A**, The Branching Point Verifier is designed as a Residual Single Head Network (RSHN), comprising a down-sampling module, an attention module, and two residual blocks. Details regarding dimension and channel down-sampling are illustrated in the inset. **B**, The Breakpoint Detector uses Residual Single Head Network (RDHN), which is similar as RSHN with the distinction of accepting two inputs: an image patch extracted around potential breakpoints and a corresponding mask image derived from the reconstruction.

**Supplementary Fig. 10.**
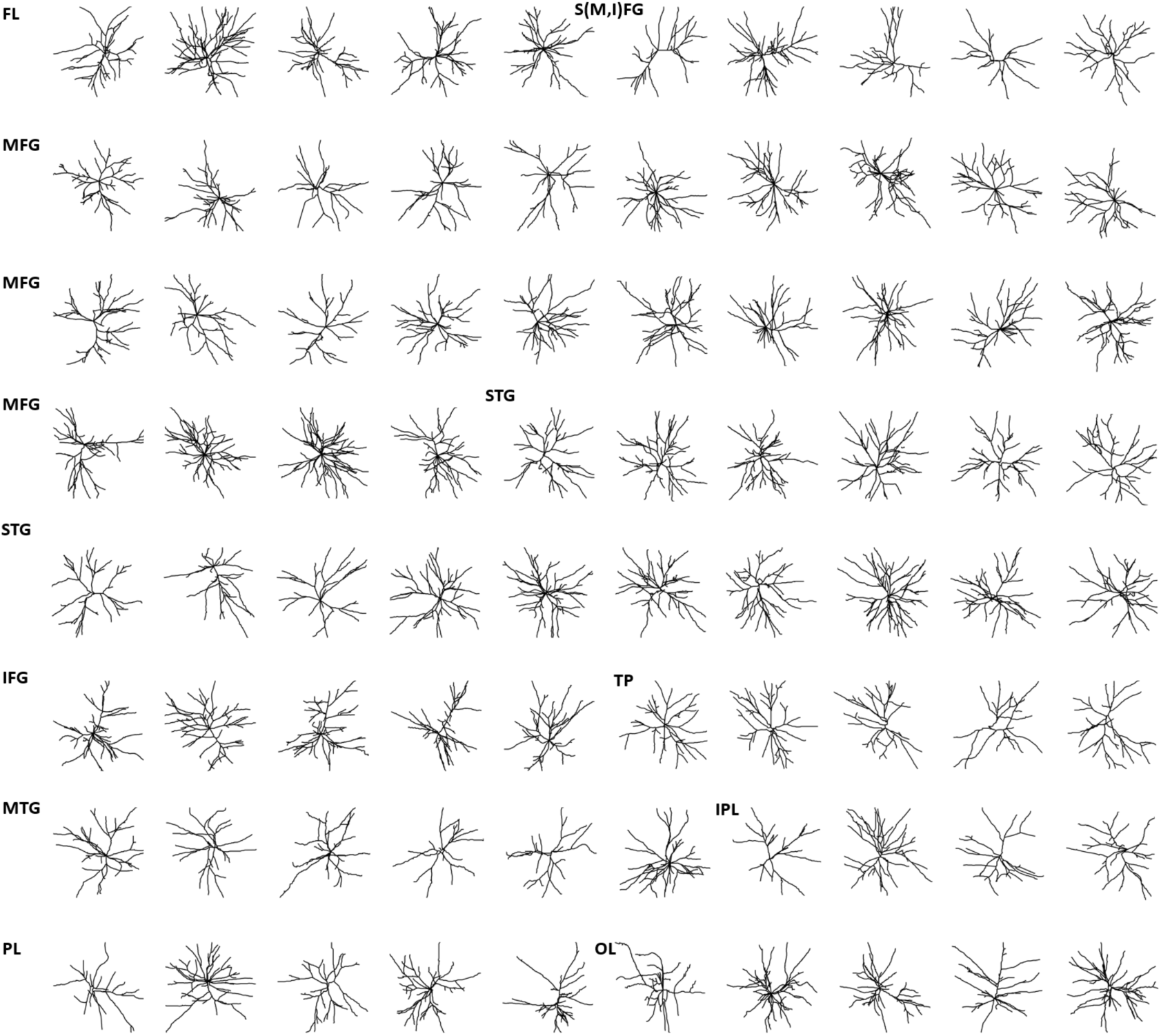
Display of 80 human neurons reconstructed using CAR. These neurons originate from 10 distinct brain regions, specifically S(M,I)FG, FL, MFG, IFG, STG, MTG, TP, PL, IPL, and OL. Region-names can be found in **Methods**.

**Supplementary Fig. 11.**
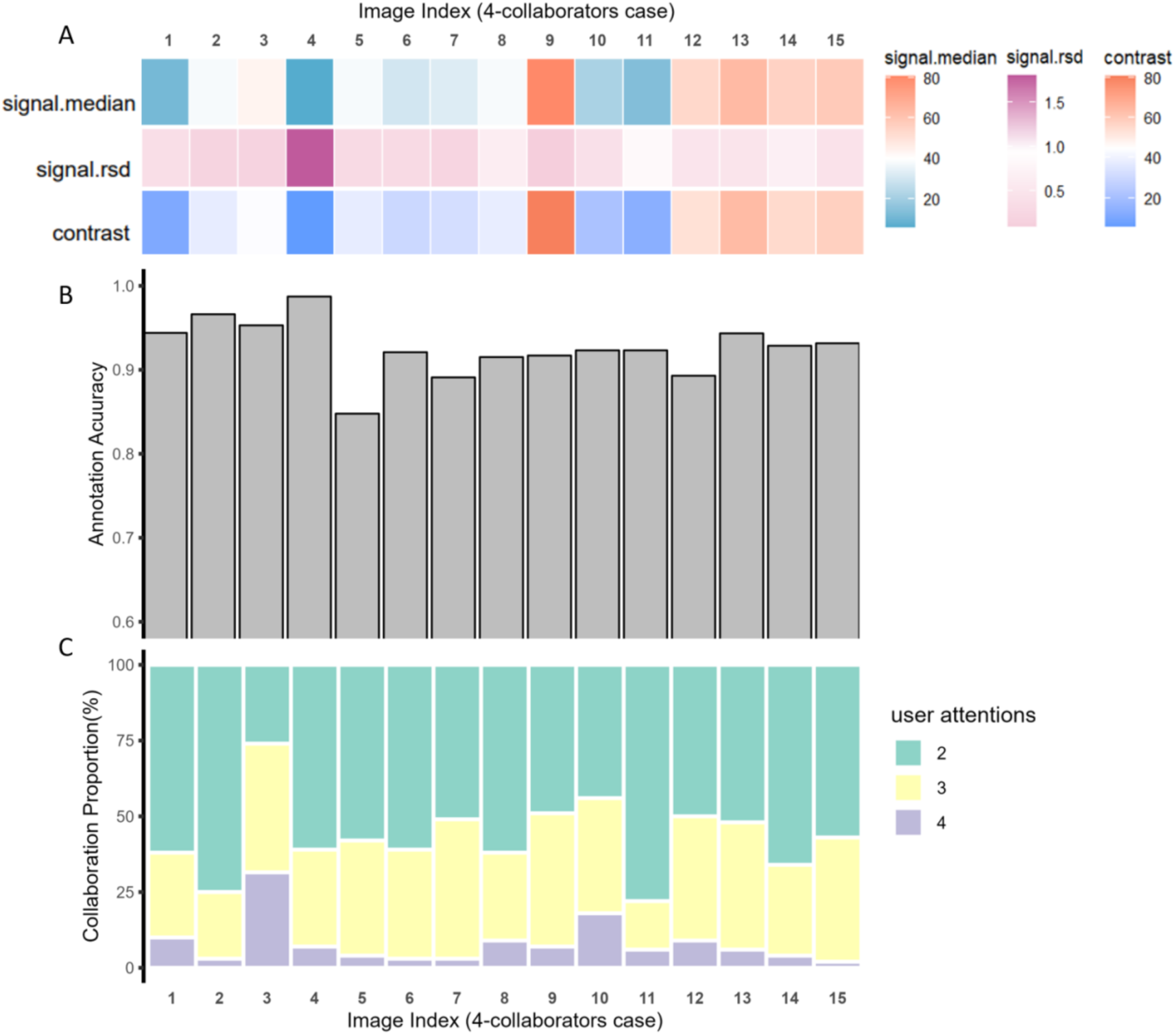
Assessment of image quality, reconstruction accuracy, and user attentions for neurons reconstructed by 4-collaborator groups. **A**, Characteristics of the 15 individual image data used by 4-collaborator groups. First row (signal.median): the median intensity of the signal (foreground). Second row (signal.rsd): the relative standard deviation (rsd) of the signal (foreground). Third row: the contrast of the image (**Methods**). **B**, The reconstruction accuracy of the 15 neurons. **C**, The degree of collaboration is illustrated using color coding: green represents the proportion of reconstructions conducted by 2 collaborators, yellow signifies 3 collaborators, and purple denotes 4 collaborators.

